# Rat Anterior Cingulate Cortex Continuously Signals Decision Variables in a Patch Foraging Task

**DOI:** 10.1101/2021.06.07.447464

**Authors:** Gary A. Kane, Morgan H. James, Amitai Shenhav, Nathaniel D. Daw, Jonathan D. Cohen, Gary Aston-Jones

**Affiliations:** Department of Psychology and Neuroscience Institute, Princeton University, Princeton, NJ; Center for Systems Neuroscience, Boston University, Boston, MA; Brain Health Institute, Rutgers University, Piscataway, NJ; Department of Cognitive, Linguistic, & Psychological Sciences and Carney Institute for Brain Science, Brown University, Providence, RI

**Author notes:** Corresponding Author: Gary Kane.

**Keywords:** Foraging, Decision-Making, Anterior Cingulate Cortex

## Abstract

In patch foraging tasks, animals must decide whether to remain with a depleting resource or to leave it in search of a potentially better source of reward. In such tasks, animals consistently follow the general predictions of optimal foraging theory (the Marginal Value Theorem; MVT): to leave a patch when the reward rate in the current patch depletes to the average reward rate across patches. Prior studies implicate an important role for the anterior cingulate cortex (ACC) in foraging decisions based on MVT: within single trials, ACC activity increases immediately preceding foraging decisions, and across trials, these dynamics are modulated as the value of staying in the patch depletes to the average reward rate. Here, we test whether these activity patterns reflect dynamic encoding of decision-variables and whether these signals are directly involved in decision-making or serve a more general function such as monitoring task performance or allocating cognitive control. We developed a leaky accumulator model based on the MVT that generates estimates of decision variables within and across trials, and tested model predictions against ACC activity recorded from rats performing a patch foraging task. Model predicted changes in MVT decision variables closely matched rat ACC activity. Next, we pharmacologically inactivated ACC to test the contribution of these signals to decision-making. Despite ACC inactivation, rats still followed the MVT decision rule, suggesting that foraging decision variables represented in the ACC are used for a more general function such as regulating cognitive control or motivation.

**Significance:** The ability to make adaptive patch-foraging decisions – to remain with a depleting resource or search for better alternatives – is critical to animal well-being. Previous studies have found that anterior cingulate cortex (ACC) activity is modulated at different points in the foraging decision process, raising questions about whether the ACC guides ongoing decisions or serves a more general purpose of regulating cognitive control. To investigate the function of the ACC in foraging, the present study developed a dynamic model of behavior and neural activity, and tested model predictions using recordings and inactivation of ACC. Findings revealed that ACC continuously signals decision variables but that these signals are more likely used to regulate cognitive control than to guide ongoing decisions.

## Introduction

Animals frequently encounter patch-foraging decisions; that is, decisions about whether to persist in harvesting a depleting resource within a patch, or to leave the patch, incurring a cost of time and effort, in search of a potentially better resource. The ability to make adaptive foraging decisions – choosing the appropriate time to leave a patch in order to maximize rewards or resources over time – is a critical skill. The mathematically optimal behavior in patch foraging tasks, described by the Marginal Value Theorem (MVT: Charnov, 1976), is to leave a patch when the *local* reward rate (the reward rate offered by the current patch) depletes below the level of the *global* reward rate (the average reward rate across all patches visited in the environment). Although animals sometimes deviate quantitatively from the predictions of this theory (Nonacs, 2001; Wikenheiser et al., 2013; Kane et al., 2019), behavior is generally qualitatively consistent with the idea that decisions are based on maximizing overall rewards by comparing estimates of the local reward rate with estimates of the global reward rate (Hayden et al., 2011; Constantino and Daw, 2015; Hayden, 2018).

Previous research into the neural mechanisms of foraging decisions has focused on the role of the anterior cingulate cortex (ACC). ACC activity is greater when the current offer of reward is more similar to the average of alternative options in foraging tasks (Hayden et al., 2011; Kolling et al., 2012; Shenhav et al., 2014). Single-unit recordings of ACC neurons in monkeys have also revealed that ACC neurons exhibit transient increases in activity around the time of foraging decisions (Hayden et al., 2011; Blanchard and Hayden, 2014), suggesting that the ACC plays a critical role in guiding foraging decisions.

The ACC has also been proposed to play an important role in cognitive control. Behaviors that require control (e.g. inhibition of an automatic response or choosing among strong competing inputs to achieve a goal) engage the ACC (Botvinick et al., 2001; Shenhav et al., 2013). The engagement of ACC in control-demanding tasks has been used to explain its dynamics during foraging decisions: in foraging tasks, as the local reward rate depletes to the level of the global reward rate, more cognitive control is needed to either i) override the default (automatic) response of staying in the patch to choose the non-default option of leaving the patch in search of better future opportunities (Kolling et al., 2012, 2016), or ii) to choose between two options with increasingly similar values, which, according to the MVT, are the most similar at the optimal time to leave the patch (Shenhav et al., 2014, 2016a, 2016b). Consistent with this interpretation, trial-by-trial changes in ACC activity in foraging tasks correlates with choice difficulty, a metric that is important for allocating cognitive control (Shenhav et al., 2014, 2016b). However, it is unclear whether encoding of metrics important cognitive control can explain within-trial dynamics of ACC activity.

In the present study, we tested whether within trial dynamics of ACC activity (e.g., transient increases around the time of foraging decisions) reflected continuous encoding of variables that are important for cognitive control, such as choice difficulty, or whether these dynamics guided ongoing foraging decisions. We developed an evidence accumulation model of foraging decision making, similar to Davidson and El Hady (2019), to i) compare changes in ACC activity to moment-by-moment changes in MVT-derived decision variables, such as the local reward rate, global reward rate, and choice difficulty, and ii) examine which components of the foraging decision process were affected by ACC inactivation. We report two key findings: changes in ACC activity within and across trials closely matched moment-by-moment changes in MVT-derived decision variables, and despite inactivation of ACC, rats retained sensitivity to foraging-related information (i.e. rats still followed the MVT decision rule).

## Materials and Methods

### Animals

Adult, male Long-Evans rats (Charles River, Kingston, NY; n = 22) were used. Rats were housed on a reverse 12 h/12 h light/dark cycle (lights off at 8 a.m.). All testing was conducted during the dark period. Throughout behavioral testing, rats were food restricted to maintain a weight of 85% to 90% ad-lib feeding weight and were given ad-lib access to water. All procedures were approved by the Rutgers University Institutional Animal Care and Use Committee.

### Foraging Task

The task was implemented using Med Associates operant conditioning chambers. Animals were trained and tested as in Kane et al. (2017) and Kane et al. (2019). Rats were first trained to lever press for 10% sucrose water on a fixed ratio (FR1) reinforcement schedule. Once exhibiting 100+ lever presses in a one-hour session, rats were trained on a sudden patch depletion paradigm — the lever stopped yielding reward after 4–12 lever presses — and rats learned to nose poke to reset the lever. Next rats were tested on the full foraging task.

A diagram of the foraging task is shown in Figure 1A. On a series of trials, rats had to repeatedly decide to lever press to harvest reward from the patch or to nose poke to travel to a new, full patch, incurring the cost of a time delay. At the start of each trial, a cue light above the lever and inside the nose poke turned on, indicating rats could now make a decision. The time from cues turning on until rats pressed a lever or used the nose poke was recorded as the decision time (DT). A decision to harvest from the patch (lever press) yielded reward (10% sucrose water) as soon as the rat entered the reward magazine. The next trial began after a 7 s inter-trial interval (ITI). With each consecutive harvest, rats received a smaller (exponentially diminished) volume of reward to simulate depletion from the patch. A nose poke to leave the patch caused the lever to retract for a delay of 10 s simulating the time to travel to a new patch. After the delay, the other lever extended, and rats could harvest from that now replenished patch. Replenished patches started with varying amounts of reward, depleting via the same exponential decay function (e.g., if the rat received 90 uL on one trial, they would receive 80 uL on the next trial regardless of the patch starting reward; Figure 1B). Rats were trained until they exhibited stable behavior across at least 3 days before testing sessions.

**Figure 1.**
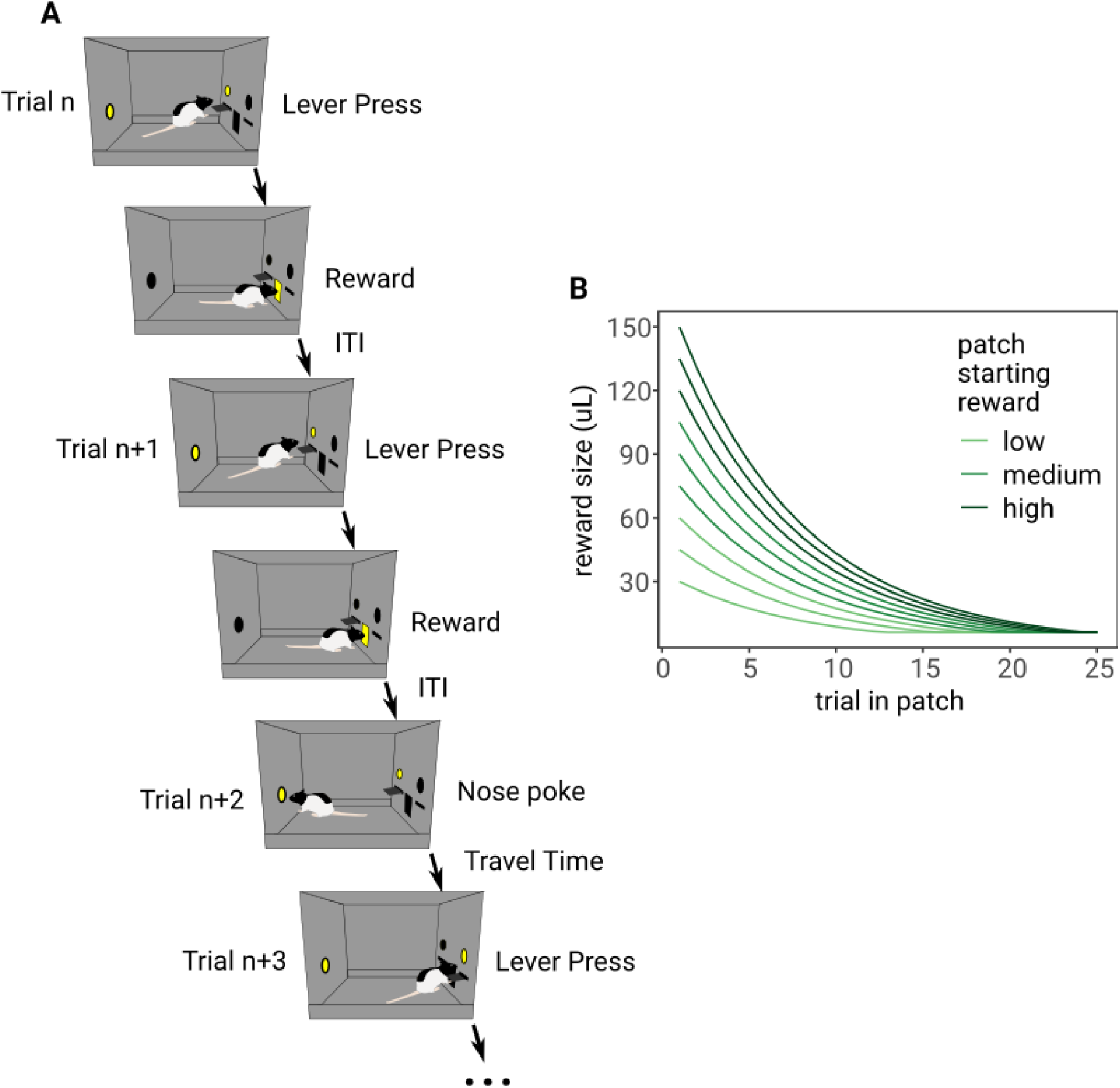
A) Operant chamber diagram of the foraging task. On trial *n*, the rat chose to press the lever to harvest from the patch, then received reward in the reward magazine in the center of the chamber. After an ITI (7 s), the rat chose to press the same lever on trial *n+1* to harvest a smaller volume of reward. On trial *n+2*, the rat chose to nose poke in the back of the chamber, initiating a “travel time” delay (10 s), after which, the rat could continue to harvest in a replenished patch by pressing the lever on the other side of the chamber (trial *n+3*). B) Reward depletion curves for each of the 9 patch starting reward volumes. Colors indicate whether the patch was a subjective low, medium, or high reward patch, for consistency with further analyses.

### Leaky Competing Accumulator Model

The model of the foraging task had two layers. The first layer, termed the value layer, consisted of two leaky accumulator units: one encoded the value of staying in the patch as the local reward rate and the other encoded the value of leaving the patch as the global reward rate. Importantly, these units were not in competition with one another (no mutual inhibition between them). The second layer, termed the decision layer, was a two-unit leaky competing accumulator layer (LCA; Usher and McClelland, 2001). The two units in this layer accumulated input from the value of staying and value of leaving units in the value layer, respectively. Additionally, there was mutual inhibition between these units. Decisions to stay vs. leave the patch on each trial were made when the activity of one of the decision units crossed a pre-defined threshold.

The value layer estimated the local reward rate (*localRate*) and the global reward rate unit (*globalRate*) by integrating reward input, *r*, at different timescales: *localRate* integrated rewards quickly but decayed quickly, and *globalrate* integrated rewards slowly but decayed slowly. The change in *localRate* and *globalRate* over time were:

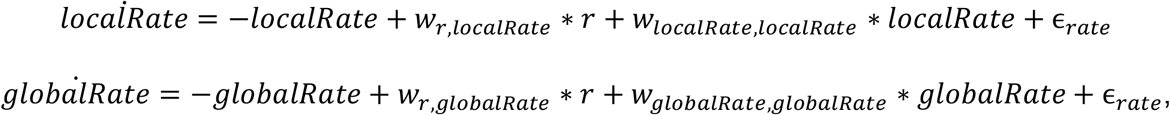

where *w_x_*_1*,x*2,_ indicates the weight between units *x*_1_ and *x*_2_ and 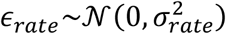. *w*_(*r*,*localRate*)_ = 1 for all simulations. *w*_*localRate*,*localRate*_, *w*_*r*,*globalRate*_, *w*_*globalRate*,*globalRate*_, and *σ*_*rate*_ were all free parameters (the noise terms for both value layer units had the same variance).

The *localRate* and *globalRate* units in the value layer units were inputs to respective units in the the decision layer, *stayDecision* and *leaveDecision*. During the decision period –– between the start of the trial and the execution of the lever press or nose poke –– decision layer units integrated input from the value layer. The activity of the decision units was:

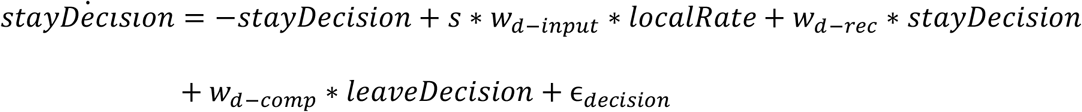

*leaveDecision*

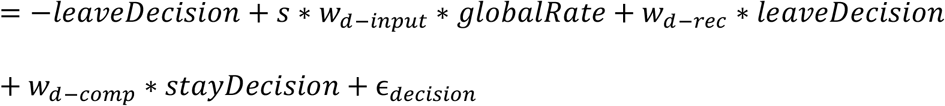

where *s* = 1 during the decision period and *s* = 0 otherwise, *w*_*d*−*input*_ is the weight between the value layer units and their respective decision layer unit (*w*_*d*−*input*_ > 0), *w*_*d*−*rec*_ is the weight of the recurrent connections in the decision layer (0 < *w*_*d*−*rec*_ < 1), and *w*_*d*−*comp*_ represents the competition between the decision units (*w*_*d*−*comp*_ < 0), and 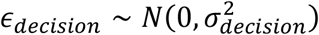. A sigmoidal activation function was used to normalize the activity of the decision units, instead of the ReLU function often used with LCA models (Usher and McClelland, 2001):

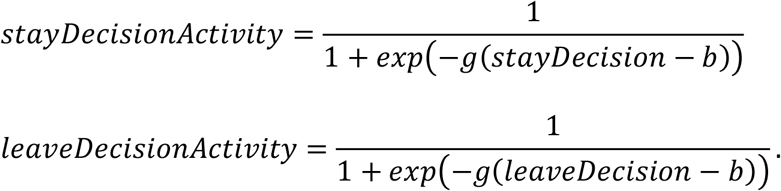

A decision to stay vs. leave was made when the activity of one of the decision units, *stayDecisionActivity* or *leaveDecisionActivity*, crossed a threshold *z*, where 0 < *z* < 1. Response times (RTs) were recorded as the time from the start of a trial until threshold crossing, plus some non-decision time. Rats’ RTs were highly variable, with many very short (< 0.1 s) responses, as well as very long responses (> 5 s). To accommodate this variability, the non-decision time was drawn from a long-tailed Weibull distribution, characterized by a mean *η* and coefficient of variation *γ*. Thus, the accumulation to bound process consisted of 9 free parameters: *w*_*d*−*input*_, *w*_*d*−*rec*_, *w*_*d*−*comp*_, *σ*_*decision*_, *z*, *g*, *b*, *η*, and *γ*.

Following decisions to stay in the patch, an additional delay (0.4 s) was added to the model to simulate the time it took rats to enter the reward port after a lever press. To model slow delivery of reward (sucrose water) from a syringe pump and the extra time rats spend consuming the reward, reward input was switched from an off state (*r* = 0) to an on state (*r* = 1) for double the duration that the syringe pump was turned on. As in the rat foraging task, the model experienced a 7 s ITI starting at the beginning of reward delivery, after which, the next trial began. Following decisions to leave the patch, the model experienced a 10 s travel time delay with no input, after which, the next trial began. The model was simulated at time steps of 0.1 s (10 steps/second).

Next, the model was fit rats’ choices and RTs. As there was no closed-form solution for the likelihood of choices and RTs, we devised a method to approximate the likelihood of choices and RTs as a function of the number of trials spent in patches and the patch starting reward volume. This method is outlined below:

1. <u>For a given set of parameters, simulate a session of the foraging task and record choices and RTs.</u> For each simulation, we ran the equivalent of a 6 hr simulation to a large sample of simulated trials. Because the global reward rate was initialized to a value of zero, the first 1 hr of the simulated choices and RTs were discarded to allow the model sufficient time to “learn” the global reward rate through experience.
2. <u>Measure the likelihood of choices to stay vs. leave as a function of the patch starting reward and the number of trials spent in the patch</u>. Simulated choices were fit with a logistic regression model using the *glm* function in R. The probability of observed choices as a function of the simulated choices was calculated using the coefficients from this logistic regression (i.e. using the *predict* function). In pseudocode:

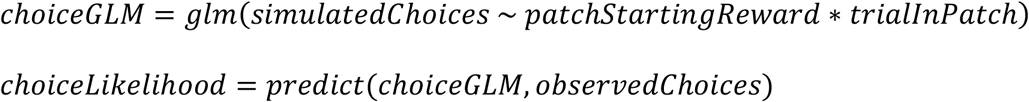

3. <u>Measure the likelihood of RTs as a function of the choice (stay vs. leave), patch starting reward, and number of trials spent in the patch</u>. Simulated RTs were fit with a linear regression model using the *lm* function in R. Coefficients from this regression model were used to predict the observed (i.e. rats’) RTs. The probability of an observed response time was assumed to be normally distributed, where the mean was equal to the predicted RT and the variance was the residual variance from the regression model. In pseudocode:

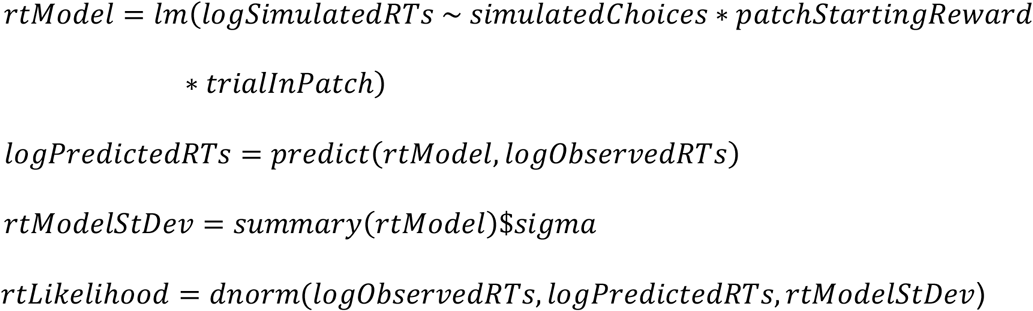

4. <u>Calculate negative log likelihood of the joint likelihood of choices and RTs:</u>

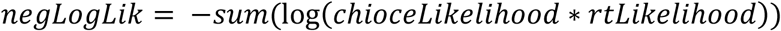

We found that this method produced a better fit to rat behavior than other approaches to fit parameters to simulated response time distributions, such as minimizing chi-square between simulated and observed response time distributions often used with diffusion models of decision-making (Ratcliff and Tuerlinckx, 2002). The maximum likelihood estimate (i.e. the parameters that minimized the negative log likelihood) was found using a genetic algorithm (the GA package in R; Scrucca, 2013).

### Electrophysiology

Prior to behavioral training, 11 rats underwent surgery to implant electrode arrays consisting of 32, 50 µm diameter single stainless-steel wires or 8 tetrodes, each consisting of 4, 25 µm diameter stainless steel wires. Wires were connected to a 32-channel Omnetics connector, serving as the interface between microwires and the headstage. First, two 0-80 machine screws were inserted into the skull over the posterior parietal cortex (approx. -4 mm AP, +/- 3 mm ML from bregma). A ground wire (125 µm stainless steel with insulation removed from 3 mm of the tip of the wire) was fixed to one of the skull screws. Next, a 4 x 2 mm craniometry was made above the anterior cingulate (Cg1), from +4 mm to 0 mm anterior-posterior from bregma and 0 mm to +/- 2 mm medial-lateral from bregma. Arrays (approximately 2 x 1 mm) were centered at approximately +2 mm anterior-posterior and positioned such that the most medial wires were just lateral to the sagittal sinus (centered at approximately 0.6 mm ML), then lowered slowly (0.1 mm / minute) down to 1.25 mm below brain surface. Once arrays were in their final position, craniotomies were filled with kwik-cast (WPI; https://www.wpiinc.com/kwik-cast-kwik-cast-sealant), then arrays were cemented to the skull using metabond (Parkell; http://www.parkell.com/c-b-metabond_3). Additional Jet Denture Repair Acrylic (Lang; https://www.langdental.com/products-Jet-Denture-Repair-Package-44) was applied over the entire surface of exposed skull and over the metabond to provide further stability to the headcap, to secure the 32-channel Omnetics connector to the skull, and this dental acrylic was shaped to provide a protective barrier in front of the microwire array and connector. After the dental cement dried, sutures were placed at the front and back of the incision as needed, and rats were returned to their home cage to recover. Rats were given meloxicam (1 mg/kg, s.c.) at the start of the surgery for analgesia and again once every 24 hr for 3 days after surgery. Rats were left to recover for one week before beginning testing.

After recovery, rats were trained 5 days per week for 3-6 weeks on the foraging task prior to recordings. One recording session was taken per rat. Prior to the recording session, a 32-channel digitizing headstage (Plexon) was plugged into the Omnetics connector on the rats ’head. From the headstage, signals were passed via a flexible cable, through a commutator, then to a Plexon Omniplex recording system. Wideband signals were sampled at 40,000 Hz. Further processing was performed in Plexon Offline Sorter software. The wideband signal for each channel was first bandpass filtered between 600-6,000 Hz and spikes were detected using a threshold of 5 times the median absolute deviation of the signal. Spike waveforms, from 1 ms before threshold crossing to 2 ms after threshold crossing, were extracted and clusters were manually identified using a combination of principal components, waveform energy, and waveform amplitude. Only clusters that exhibited consistent firing throughout the entire session were included for analysis. Clusters were characterized as single units if less than 2% of spikes within the cluster exhibited an inter-spike interval of less than 2 ms, and the cluster had an L-ratio (Schmitzer-Torbert et al., 2005) of less than 0.1. All other clusters were characterized as multi-units. Altogether, this resulted in a total of 42 single-units and 106 multi-units. All units were combined together for all further analyses.

After the completion of recordings, small electrolytic lesions were made by passing current (25 uA for 15 s) through wires at the front and back of the array. 24 hr later, rats were perfused with 4% paraformaldehyde (PFA) and their brains were extracted. Brains were post-fixed in 4% PFA for 24 hr, then cryoprotected in 30% sucrose in phosphate buffered saline for 72 hrs. Finally, brains were flash frozen and sectioned into 40 µm sections on a cryostat. Electrode locations were confirmed by locating lesions.

### Pharmacological Inactivation of ACC

Prior to behavioral training,11 rats underwent surgery to implant a bilateral cannula targeting the ACC (Cg1). Similar to electrode array implant surgeries, two 0-80 machine screws were inserted into the skull above the posterior parietal cortex. Next, a large craniotomy was drilled, spanning the ACC bilaterally (from approx. -1 to +1 mm ML and +1 to +3 mm AP from bregma). The bilateral cannula (PlasticsOne) was positioned to target Cg1 at +/- 0.5 ML and +2 mm AP from bregma. The cannula was lowered slowly (0.1 mm/minute) to a depth of 0.75 mm below the brain surface. The implant was secured to the skull using metabond, and the Jet Denture acrylic was used to further secure the implant to the skull and skull screws, and it was shaped to create a protective barrier in front of the cannula. Following completion of the surgery, sutures were applied as needed to secure the front and back of the incision and was then placed in its home cage to recover. Rats followed the same analgesia protocol and post-operative recovery as with electrode array implants.

After full recovery, rats were trained 5 days/week for 4 weeks on the foraging task before testing. On test days, 15 minutes prior to the start of the session, rats underwent a microinjection of either a cocktail of the GABA agonists baclofen and muscimol (Bac-Mus; 1 mM and 0.1 mM, respectively; 0.5 μL/side), or artificial cerebro-spinal fluid (aCSF, 0.5 μL/side) as a control. A bilateral injector (33 G, PlasticsOne) that protruded 0.5 mm below the bottom of the cannula (to a depth of 1.25 mm) was inserted through the cannula, and Bac-Mus or aCSF was injected at a rate of 100 nL/min. The injector was left in place for 2 minutes after completion of the injection to allow the drug cocktail to diffuse into the tissue and to avoid backflow of the drug cocktail up the cannula track. The injector was then removed, and rats were placed in the operant chamber awaiting testing. The day before the first injection, rats underwent one sham injection to acclimate to the procedure. Rats were tested with Bac-Mus and aCSF for one session with each drug, counterbalanced (4 rats received aCSF followed by Bac-Mus, 4 vice-versa). Rats were given one recovery day, in which they were tested without an injection, between the two testing sessions.

## Experimental Design and Statistical Analysis

### Rat foraging behavior

All statistical analyses and computational modeling were conducted in R (R Core Team, 2020). Mixed effects (ME) models were fit using the *lme4* package (Bates et al., 2015), and significance tests for linear mixed effects models were performed using the *lmerTest* package (Kuznetsova et al., 2017). Unless otherwise specified, all continuous predictors in mixed effects models were z-scored.

To investigate the behavioral performance of rats that participated in the ACC recording experiment, we analyzed rats’ foraging decisions (the number of trials spent in each patch) and response times (the time from the start of the trial until the lever press or nose poke) during the final three training sessions prior to the recording session. Looking at their training data allowed us to pool behavior across multiple sessions. In this experiment, two main hypotheses were tested: i) that rats would spend more trials in patches that offered greater rewards (a standard prediction of the MVT); and ii) as patches depleted, response times to decide to stay vs. leave would increase (reflecting greater decision difficulty). The first hypothesis was tested using a linear mixed effects model of the number of trials spent in each patch, with a fixed effect of the starting reward volume of the patch and random intercept for each rat (lme4 syntax: *TrialsInPatch* ∼ *PatchStartingReward* + *PatchStartingReward*|*Rat*)). To test whether rats adopted the same reward rate leaving threshold across patches, a mixed effects model of the reward rate at the time rats left patches, with a fixed effect of the patch starting volume and random intercept for each rat. In this model, to directly compare the leaving threshold at each of the 9 patch starting reward volumes, patch starting reward was treated as a categorical variable (dummy coded) and we conducted pairwise comparisons of the reward rate when rats left the patch across all patch starting reward volumes.

The second hypothesis, that response times would increase as patches depleted, was initially tested using a linear mixed effects model of the log of response times with fixed effects of the patch starting reward volume, the number of trials spent in the patch, and the starting reward x trials in patch interaction, and a random intercept for each rat (*logRT* ∼ *PatchStartingReward* * *TrialsInPatch* + *PatchStartingReward* * *TrialsInPatch*|*Rat*)). The log of response times was used as the raw response times were positively skewed. However, if rats exhibit longer response times as patches deplete, then it is likely that they will exhibit longer response times, on average, in patches that start with smaller rewards because a greater proportion of trials spent in these patches will be at lower reward volumes. To better examine whether there were differences in response times across patches, another linear mixed effects model was used to examine the effect of the number of trials remaining in the patch and patch starting reward on the log of response times (*logRT* ∼ *PatchStartingReward* * *exp*(*TrialsRemaining*) + (*PatchStartingReward* * *exp*(*TrialsRemaining*)|*Rat*)). An exponential function of *TrialsRemaining* was used, as it proved to be a better fit to data than a linear function (Figure 1E). In this model, if the response times around the trial at which rats chose to leave the patch were similar across different patch types, there would be no main effect of *PatchStartingReward*.

### Leaky Accumulator Model Predictions

Simulated behavioral data and the time course of activity of each of the LCA units were obtained via simulations as described above. To generate peri-event time histograms (PETHs) of LCA unit activity around the time of decisions, first, the time of decisions was obtained from the simulated behavioral data. Next, for each trial, the activity of each LCA unit was extracted for 8 simulated seconds (80 time steps) before the decision and 4 s (40 time steps) after. In addition, PETHs were created for the relative value of leaving the patch, decision difficulty, and decision conflict. The relative value of leaving the patch was the moment-by-moment difference between the local and global reward rate units: *globalRate* − *localRate*; decision difficulty or the similarity in the value of staying vs. leaving was defined as: −*abs*(*localRate* − *globalRate*); and decision conflict was the product of the activity of the decision units:

*stayDecisionActivity* * *leaveDecisionActivity*. Each of these variables were first normalized (z-scored), then PETHs were created the same as for the 4 LCA model units.

### Analysis of ACC activity

To analyze the correlation between average ACC activity and foraging decisions and response times, first, the firing rate of each unit was calculated for each trial. A linear mixed effects model were used to test the effect of trials until leaving and starting patch reward, with random effects for all parameters for each unit:

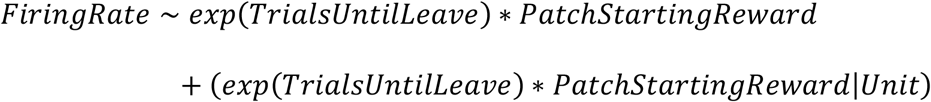

A second linear mixed effects model was used to test the effect of the log of response times, with random intercepts and slopes for each unit:

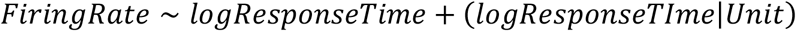

To examine i) at what point in the decision process ACC activity was influenced by foraging decisions and response times and ii) which units encoded these two variables over the course of the entire trial, PETHs were created for each unit, from 8 s before the lever press to stay in the patch to 4 s after the lever press, in time bins of 0.1 s (120 total time points). Next, 3 generalized linear models (GLMs) with a quasi-poisson link function (poisson regression with an overdispersion parameter) were fit to the spike counts in each bin of the PETH for each unit. The first model included an intercept, effect of trials until leaving, and effect of the log of response times for each time point (a total of 360 parameters). Coefficients of the effect of trials until leaving and effect of log of response times were used to determine the strength of encoding of these variables at that specific point within the trial. To determine whether these variables significantly contributed to the variance in each unit’s activity, separate GLMs that excluded one of the effect of trials until leaving or the effect of log of response times were fit to each unit, and likelihood ratio tests were conducted between the full model and the models excluding one of these variables. If the full model provided a better fit to a specific unit’s PETH, assessed via likelihood ratio test against a model with one predictor removed, that would indicate significant encoding of the variable excluded from the reduced model. Based on this analysis, units were characterized as encoding the number of trials until leaving, the log of response times, or both.

Finally, the dynamics of ACC activity within trials, and the modulation of these dynamics was qualitatively compared to moment-by-moment changes in decision variables derived from the LCA model. To compare the dynamics of the average ACC neuron, an average PETH was constructed by taking the PETH for each unit described above, normalizing the activity of each unit – taking the z-score of activity across all bins for that unit – and taking the average normalized activity within each bin across units. To examine the diversity in encoding across units, the average, normalized PETH was calculated for each unit on a subset of trials leading up to the decision to leave the patch: on 5, 3, 1, or 0 trials until leaving the patch. For each unit, these 4 PETHs were concatenated into a vector with 480 features (120 timepoints for each of the 4 PETHs), and principal components analysis (PCA) was performed on these 480 features across all 148 units to extract the dimensions that capture the most variance across all units at each time point for each of these 4 trials until leaving the patch. Principal components, representing the dimensions which captured the most variance across units on these trials, were qualitatively compared to LCA model units.

### Pharmacological inactivation of ACC

Foraging behavior in the inactivation experiment was analyzed in a similar manner as the described above. Linear mixed effects models were used to test the effect of ACC inactivation (aCSF injection vs. Bac-Mus injection) on the number of trials spent in the patch (lme4 syntax: *TrialsInPatch* ∼ *PatchStartingReward* * *Inactivation* + (*PatchStartingReward* * *Inactivation*|*Rat*)) and response times (lme4 syntax: *logRT* ∼ *exp*(*TrialsUnitLeaving*) * *PatchStartingReward* * *Inactivation* + (*exp*(*TrialsUnitLeaving*) * *PatchStartingReward* * *Inactivation*|*Rat*)). An additional mixed effects model was used to test the relative value of leaving the patch (the difference in the global and local reward rate) at the time that rats chose to leave the patch, termed the *valueATLeaving*. This analysis was designed to measure whether ACC inactivation caused rats to overharvest to a greater degree than observed in control sessions. *valueATLeaving* was calculated as follows:

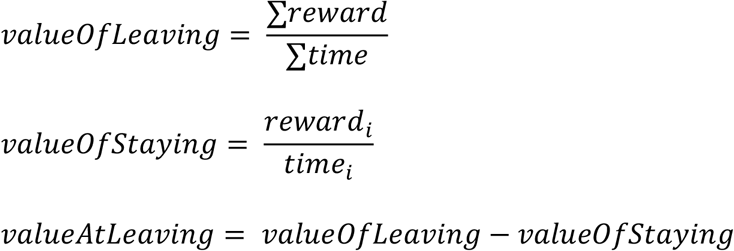

The value of leaving across the three patch types was tested as a function of the patch starting reward and drug treatment (ACC inactivation vs. control; lme4 syntax:

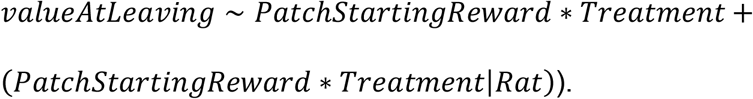

Finally, the LCA model was fit to each drug treatment as described above. Paired t-tests were run on each of the 13 parameters, and holm-corrected p-values (Holm, 1979) are reported.

## Results

### Rats spend more time in patches that offer greater rewards

Rats (n = 11) were trained to perform a patch foraging task in which they randomly encountered patches with starting rewards that ranged from 30–150 μL (Figure 1B). This wide range in reward offered by different patches tested whether rats ’ followed a central prediction of MVT: when offered greater levels of reward, rats should harvest for more trials until these patches deplete to the leaving threshold (the global reward rate or average reward rate across all patches). To test this prediction, rat behavior during their final three training sessions was analyzed. Rats participated in 553-1162 trials, visiting 99-202 patches each. As in a previous study (Kane et al., 2017), rats harvested for more trials in patches that started with greater rewards (main effect of patch starting reward: β = 2.294, SE = 0.079, F(1, 11.332) = 837.14, p < .001; Figure 2A). Among patches that started with greater rewards (75-150 μL), there was no difference in the reward rate at which rats chose to leave patches. However, rats left patches that started with smaller rewards (30-60 μL) at a lower reward rate than patches that started with greater rewards (Figure 2B, pairwise chi-square tests presented in Figure 2-1). As predicted by MVT, rats adopted a constant reward rate threshold at which to leave patches when in patches that yielded larger rewards, but contrary to MVT, they exhibited a bias to harvest reward beyond this threshold in patches that yielded smaller amounts of reward.

**Figure 2.**
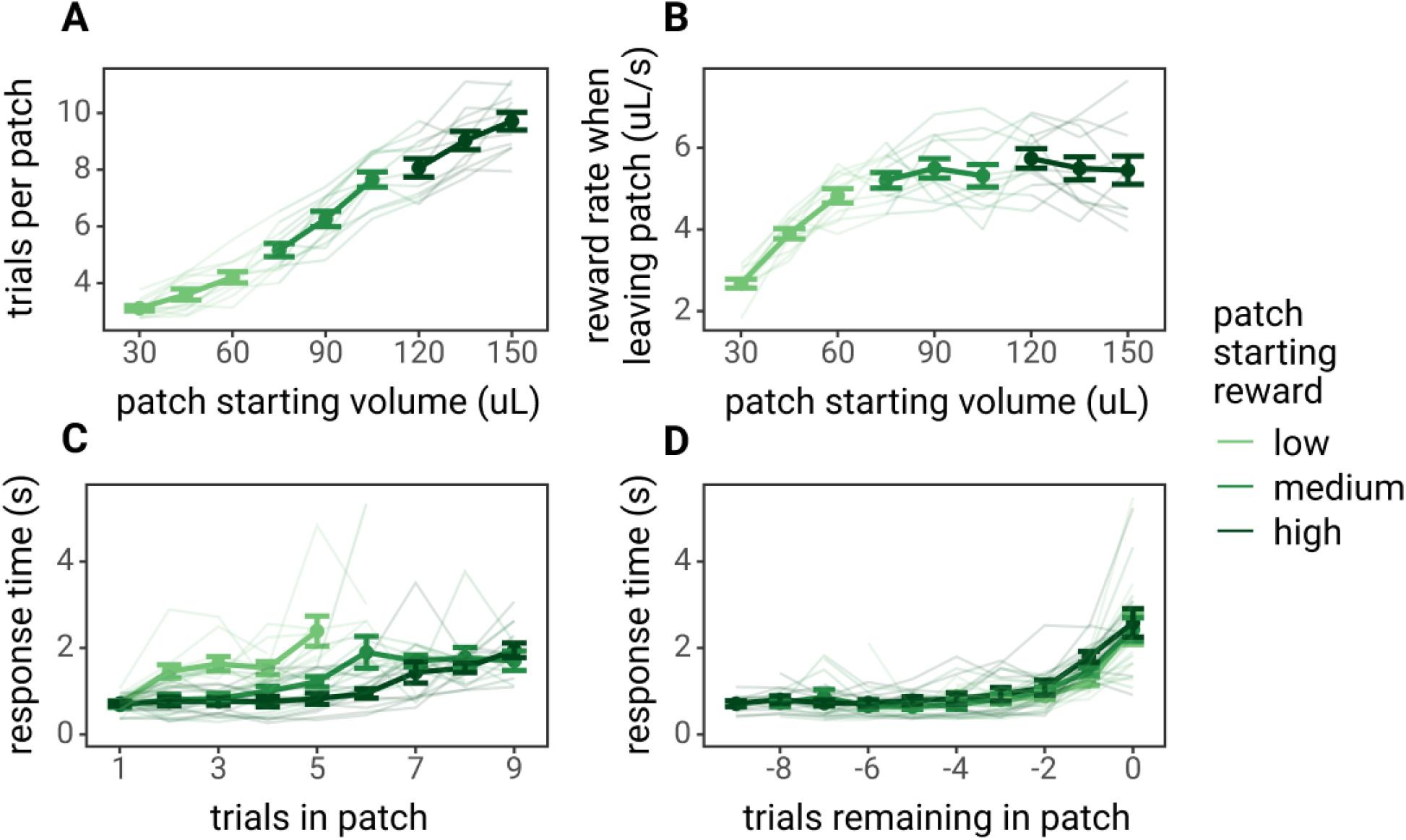
Rat behavior in the patch foraging task. In all panels, points, dark lines, and error bars represent the mean and standard error across rats, and lighter lines represent behavior of each individual rat. A) The average number of trials spent in each patch as a function of the starting reward volume of the patch. B) The average reward rate (reward volume / trial time) rats received on the trial before rats chose to nose poke to leave the patch. C) The average of the median response times for each rat over the course of trials in the patch, split by patches that started with low (30-60 μL), medium (75-105 μL), or high (120-150 μL) starting reward volumes. D) The average of the median response times for each rat as rats became closer to leaving patches. 0 trials remaining in the patch indicated the trial in which they nose poked to leave the patch, -1 trial remaining is the last lever press to stay in the patch, -2 is the second to last lever press before leaving the patch, etc.

### Rats’ response times increase as patches deplete

As patches depleted, animals may have experienced either an increased need to override the default response of choosing to stay in the patch in favor of leaving the patch or increased difficulty to decide to stay vs. leave as the value of staying vs. leaving became more similar. If rats experienced an increased need to override the default response or increased decision difficulty as patches depleted, their response times (RTs) should have increased. As rats spent more time in patches, their response times increased (main effect of trials in patch: β = 0.512, SE = 0.056, F(1, 9.791) = 85.203, p < .001; Figure 2C). Furthermore, their response times were, on average, greater in lower rewarding patches (β = -0.398, SE = 0.046, F(1, 9.830) = 75.67, p < .001). Slower average response times in lower starting reward patches is likely due to a greater proportion of trials spent with lower reward volumes – as lower starting reward patches start out in a depleted state, there are few to no trials in which rats should exhibit faster response times to stay in the patch. To test this hypothesis, response times were also analyzed as a function of the number of trials remaining in the patch. If rats experienced a reduced (or increased) need to override a default response or decision difficulty in patches that started with smaller rewards, then response times should have become faster (or slower) as rats approached the point to leave these patches. Across all patch types, rats’ response times increased as they approached the point at which they left patches (main effect of trials remaining in the patch: β = 0.555, SE = 0.058, F(1, 10.751) = 92.662, p < .001; Figure 2D), but there was no difference in the average response times (main effect of patch starting reward: β = 0.691, SE = 0.118, F(1, 9.963) = 0.002, p = 0.968) or in the rate at which response times increased among different patch types (trials remaining x patch starting reward interaction: β = 0.010, SE = 0.018, F(1, 9.663) = 0.290, p = .602; Figure 2D). Despite leaving smaller starting reward patches at a lower threshold than higher starting reward patches, rats exhibited similar response times in these patches as they became closer to leaving, suggesting that they experienced the same need to override the default response to stay in the patch or the same decision difficulty when deciding to leave patches that started with greater rewards. Alternatively, this increase in response time as patches depleted could be interpreted as a reduction in motivation or response vigor in anticipation of smaller rewards.

### A leaky competing accumulator (LCA) model of rat foraging behavior

Evidence accumulation models have proven successful in describing not only perceptual decisions requiring moment-to-moment sampling of sensory information, but also value-based decisions (Polanía et al., 2014; Tajima et al., 2016; Pisauro et al., 2017; Frömer et al., 2019; Lin et al., 2020; Peters and D’Esposito, 2020; Callaway et al., 2021). Recent theoretical work has applied the evidence accumulation framework to foraging decisions (Davidson and El Hady, 2019). To describe rats’ foraging decisions as a function of their moment-by-moment estimate of the local vs. global reward rates, we developed an evidence accumulation model that implemented the MVT decision rule – to leave a patch when the local reward rate in the current patch depletes to the level of the global reward rate – using leaky accumulators. The model consisted of two layers, the value and decision layer. The value layer units estimated the local and global reward rate by integrating rewards at different timescales: the local reward rate unit integrated rewards quickly but decayed quickly, whereas the global reward rate unit integrated rewards slowly but decayed slowly. These units were not in competition with one another – there was no reciprocal inhibition between them. The decision layer was an LCA (Usher and McClelland, 2001) that implemented an accumulation to bound process. At the start of the trial, decision layer units integrated the activity of the value layer units until one of the decision units crossed a threshold, at which point, the model chose the corresponding option (Figure 3A). A demonstration of the activity of the value layer and decision layer units during a simulation is shown in Figure 3B-C. The model was fit to rats’ choices and response times (see Materials and Methods for details, fit parameter estimates in Figure 3-1). This LCA model, fit to rat behavioral data, captured important features of rats’ behavior: the model predicted spending more trials in patches that yielded larger rewards and predicted longer response times as patches depleted, with longer response times for patches that started with smaller rewards (Figure 3D-E).

**Figure 3.**
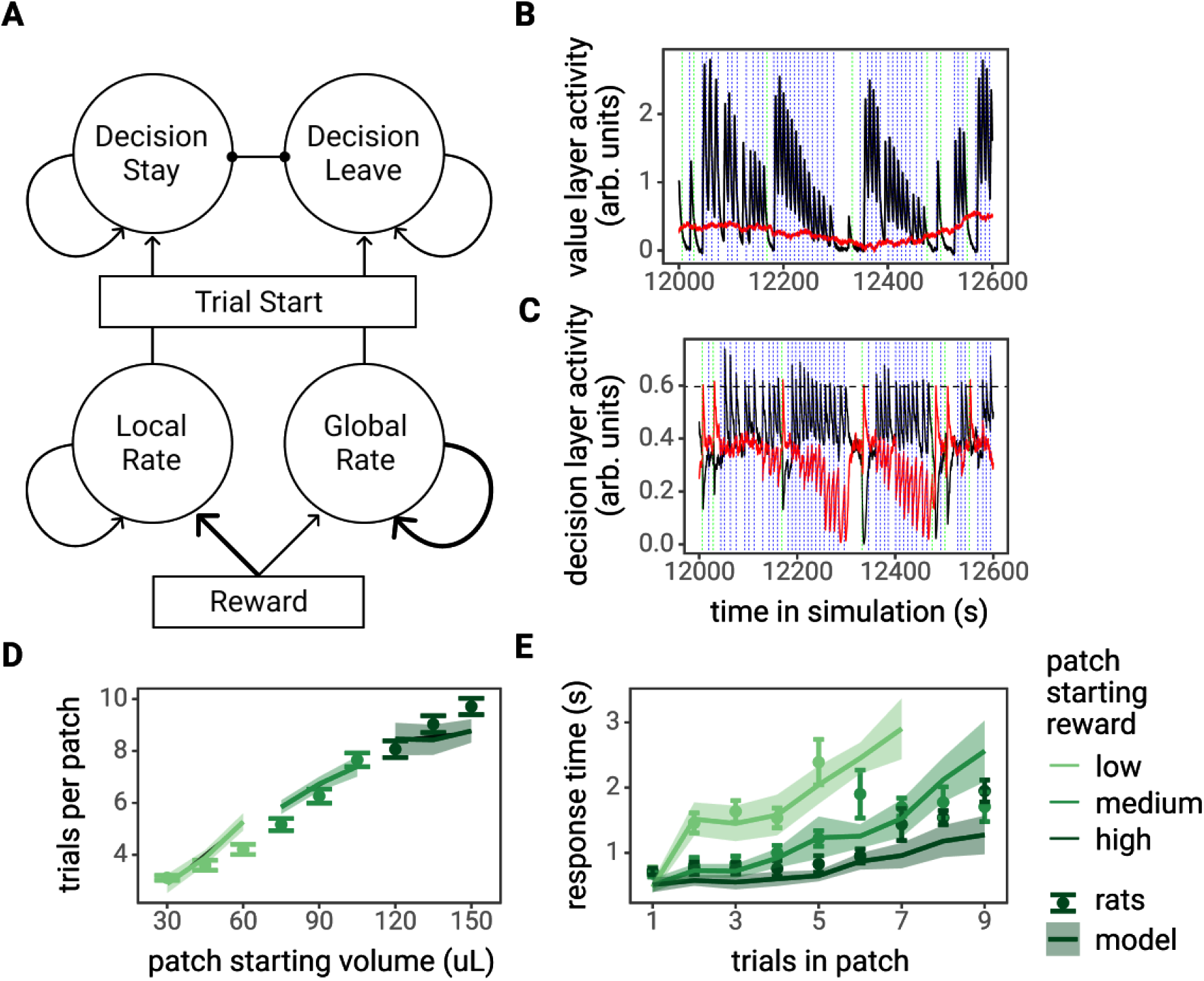
The leaky accumulator model of the foraging task. A) Diagram of the model, consisting of two layers of leaky accumulators. The bottom layer, the value layer (“local rate” and “global rate” units), estimated the local reward rate and global reward rate by integrating over rewards on different timescales. The top layer, the decision layer, was a leaky competing accumulator model that made decisions to stay vs. leave via an accumulation to bound process at the start of the trial, with input from the local and global rate units. B-C) Example activity of the value layer and decision layer units during a 10 min (600 s) sample of a model simulation. The solid black lines represent the local reward rate and decision stay unit activity, and the solid red lines represent the global reward rate and decision leave unit activity. The dotted blue and green vertical lines indicate the start of a trial in which the model decided to stay in the patch or to leave the patch, respectively. The horizontal dashed line in C represents the decision threshold. D-E) The leaky accumulator model-predicted number of trials spent in each patch type (D) and predicted response times as by the number of trials spent in patches (E) plotted against observed rat behavior. Points and error bars represent the mean +/- standard error across rats, lines and ribbon represents the mean +/- standard error of model

The LCA model was then used to generate predictions regarding ACC activity in the foraging task. As the model estimates important MVT decision variables – the local and global reward rates – on a moment-by-moment basis, the activity of LCA model units was used to calculate specific decision variables that the ACC has been hypothesized to encode. Three particular hypotheses were tested: i) ACC encodes the relative value of leaving a patch as the difference between the global reward rate and local reward rate (Kolling et al., 2012), ii) ACC encodes decision difficulty or the similarity in the value of staying and the value of leaving a patch (Shenhav et al., 2014), and iii) ACC encodes the conflict between choosing to stay vs. choosing to leave a patch, defined as the product of the decision units (Botvinick et al., 2001). The relative value of leaving the patch and decision difficulty hypotheses are equivalent while the rat is in the patch, however, they differ during the travel time. Peri-event time histograms (PETHs) of the activity of each model unit, as well as the relative value of leaving the patch, decision difficulty, and decision conflict, were created from simulation data by averaging the value of these variables over trials, locked to the time of the decision (time of execution of the lever press or nose poke in the simulation; Figure 3-2). These PETHs were later compared to recorded ACC activity.

### ACC activity correlates with foraging decisions and response times

First, we examined whether changes in ACC activity (n = 148 units, Figure 4-1) correlated with rats’ foraging decisions and response times. Consistent with the hypothesis that ACC activity should increase with the increased need for to override a default response or with increased decision difficulty as patches deplete, average ACC activity over the course of trials (the number of spikes during trial / time of trial, averaged across units) increased as rats became closer to leaving a patch (main effect of trials until leave: *β* = 0.287, SE = 0.061, F(1, 147) = 22.028, p < .001). Similar to the relationship between trials until leaving the patch and response times, average ACC activity as rats became closer to leaving a patch was not influenced by the patch starting reward (main effect of patch starting reward: *β* = 0.008, SE = 0.043, F(1, 138) = 0.032, p = 0.858). And the rate at which ACC activity increased as rats became closer to leaving a patch did not depend on the patch starting reward (patch starting reward x trials until leave interaction: *β* = 0.013, SE = 0.019, F(1, 145) = 0.509, p = 0.477; Figure 4A). Accordingly, average ACC activity over the course of trials increased linearly with response times (main effect of response times: *β* = 0.175, SE = 0.038, F(1, 148) = 20.883, p < .001; Figure 4B). No differences were noted between single- and multi-units (Figure 4-2A, 4-2B).

**Figure 4.**
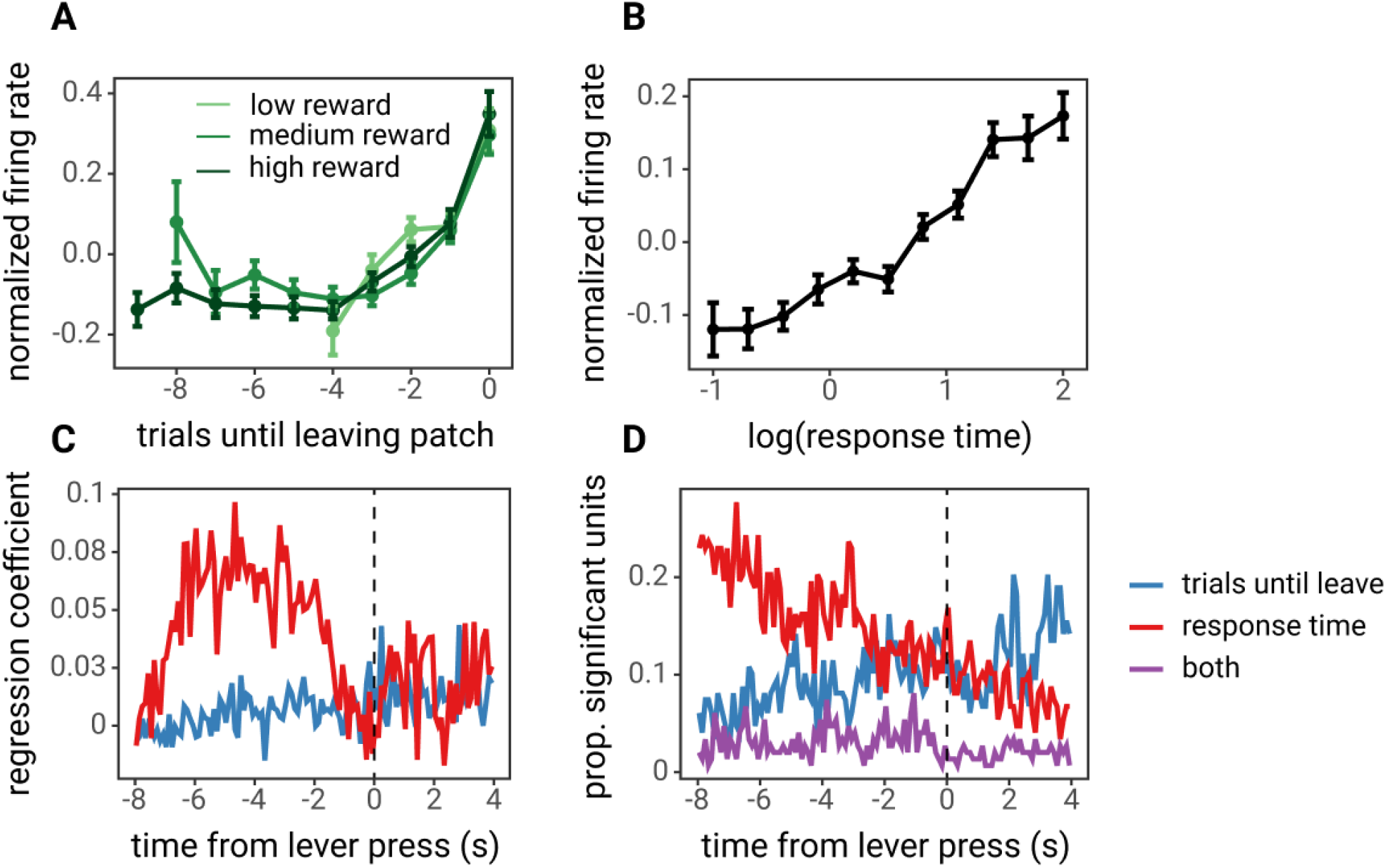
ACC activity correlates with foraging decisions and response times. A-B) Average ACC activity over the course of entire trials, normalized and averaged across units, as a function of A) trials until leaving the patch and the patch starting reward and B) the log of response times. Points and lines represent the mean normalized (z-scored) activity across units and error bars represent the standard error across units. C) The average effect of trials until leaving and response times at each time point within a trial, locked to the time of the lever press to stay in the patch. D) The proportion of units with significant effects of trials until leaving, response times or both (p < .05, z-test on regression coefficient), at each time point within the trial.

To investigate the effect of rats’ foraging decisions and response times on ACC activity in more detail, PETHs of activity around the time of the lever press to stay in the patch were created for each unit, and a series of generalized linear models (GLMs) was used to examine i) at which point in the decision process ACC encoded the number of trials until leaving the patch vs. the response time on a given trial, and ii) which units significantly encoded either variable over the entire course of the trial (see Methods for full details). Encoding of the number of trials until leaving the patch was weakest during the time period before decisions and grew stronger after decisions (through the reward and inter-trial interval periods), evidenced by increasing average regression coefficients and an increase in the number of units with a statistically significant regression coefficient for the number of trials until leaving (with p < .05, z-test; Figure 4C-D). At the same time, encoding of response times was strongest preceding decisions, with the strongest regression coefficients occurring during a window of ∼6 s to 2 s preceding the decision, and a greater number of units encoding response times preceding the decision vs. after the decision (Figure 4C-D). Lastly, we found that a large number of units (69 of 148) encoded both response times and trials until leaving – excluding one of these variables resulted in a worse model fit according to likelihood ratio tests (p < .05 with holm correction for multiple comparisons across 148 units) – with additional units encoding either response times only (36 / 148) or trials until leaving only (6 / 148). Again, no differences in encoding were observed between single- and multi-units (Figure 4-2C, 4-2D).

### ACC activity continuously tracks decision variables

To further examine what was driving encoding of response times prior to the decision and decisions later in the trial, PETHs of recorded ACC activity were compared to decision variables derived from the LCA model. First, we discovered that average normalized ACC activity – the average PETH across neurons, split by the number of trials until leaving the patch – closely tracked decision difficulty or the similarity in the value of staying in the patch vs. leaving the patch (the relative difference between the *localRate* and *globalRate* units; Figure 5A). Importantly, both decision difficulty and average normalized ACC activity i) increased leading up to decisions, ii) was inhibited during reward delivery following a decision to stay in the patch (5, 3, or 1 trial until leaving), and iii) following decisions to leave (0 trials until leaving), maintained elevated activity. This finding indicates that the dynamics of ACC activity within and across trials are consistent with the hypothesis that they encode decision difficulty.

**Figure 5.**
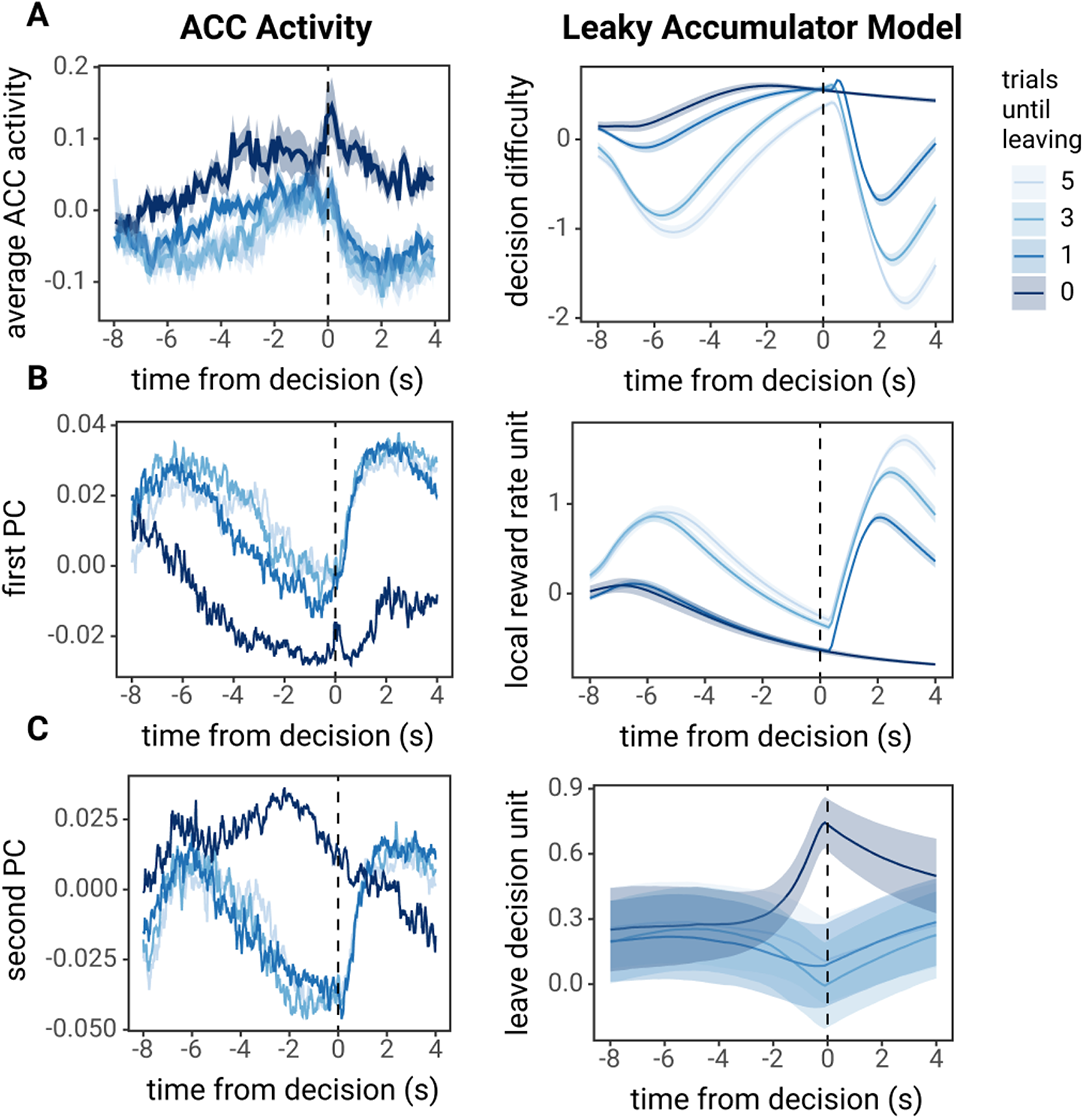
ACC activity next to decision variables derived from the leaky accumulator model. A) Average, normalized PETH across units for 5, 3, 1, or 0 trials until leaving next to decision difficulty (relative difference between the local and global reward rate). B-C) PCA was performed across the PETH for all units on these trials (5, 3, 1, and 0 trials until leaving). B) The first PC next to the local reward rate unit activity. C) The second PC next to the leave decision unit activity. For leaky accumulator model unit activity, PETHs of decision variables were created from simulations using parameters fit to each rat, Lines and ribbon represent the average across these simulations.

To better understand the heterogeneity in the modulation of ACC firing across different units, we created PETHs for each unit (both single- and multi-units) at 5, 3, 1, and 0 trials until leaving the patch, and performed principal components analysis (PCA) on these PETHs for each unit (including 120 time points x 4 trials = 480 features and 148 units or observations). Next, we compared the principal components (PCs) – the dimensions which explained the most variance in ACC activity (Figure 5-1) – to decision variables derived from the LCA model. Whereas the average PETH across all units closely tracked decision difficulty, the first two PCs closely tracked different LCA model units. The first PC exhibited activity similar to the local reward rate, *localRate*, with i) decreasing activity leading up to decisions, ii) a sharp increase in when receiving reward (immediately after the decisions to stay), and iii) consistently reduced activity on decisions to leave. The second PC was similar to the leave decision unit, *leaveDecisionActivity*, with activity that i) decreases leading up to decisions to stay and ii) increases leading up to decisions to leave. These findings suggest that while on average, ACC signals decision difficulty, this signal is made up of units that signal lower-level decision variables, including reward rate and decision accumulators.

### ACC inactivation does not alter the foraging decision process

Continuous encoding of decision variables in ACC could play a central role in decision-making or indicate a more general role such as monitoring ongoing performance for the purpose of allocating cognitive control to decisions. To test the contribution of the ACC to foraging decisions, ACC was pharmacologically inactivated via microinjection of a cocktail of the GABA receptor agonists baclofen and muscimol (Bac-Mus) immediately prior to testing rats in the foraging task. In this experiment, rats were tested on a simplified version of the foraging task, with only three starting patch reward volumes (60, 90, and 120 μL). Compared to control sessions in which rats were injected with artificial cerebrospinal fluid (aCSF), ACC inactivation caused rats to stay in patches for more trials (main effect of aCSF vs. Bac-Mus: *β* = 2.397, SE = 0.227, F(1, 870) = 111.913, p < .001; Figure 6A), and increased response times as rats came closer to leaving the patch (main effect of aCSF vs. Bac-Mus: *β* = 0.669, SE = 0.054, F(1, 7876) = 149.849, p < .001; Figure 6B). However, despite ACC inactivation, rats still stayed longer in patches that started with greater rewards (main effect of patch starting reward: *β* = 1.635, SE = 0.144, F(1, 869) = 202.462, p < .001; no patch starting reward x treatment interaction; *β* = 0.056, SE = 0.226, F(1, 869) = 0.062, p = 0.803), and rats still exhibited longer response times as they became closer to leaving the patch (main effect of trials until leaving: *β* = 0.287, SE = 0.022, F(1, 7876) = 277.213, p < .001; no trials until leaving x treatment interaction; *β* = 0.043, SE = 0.032, F(1, 7877) = 1.876, p = .171).

**Figure 6.**
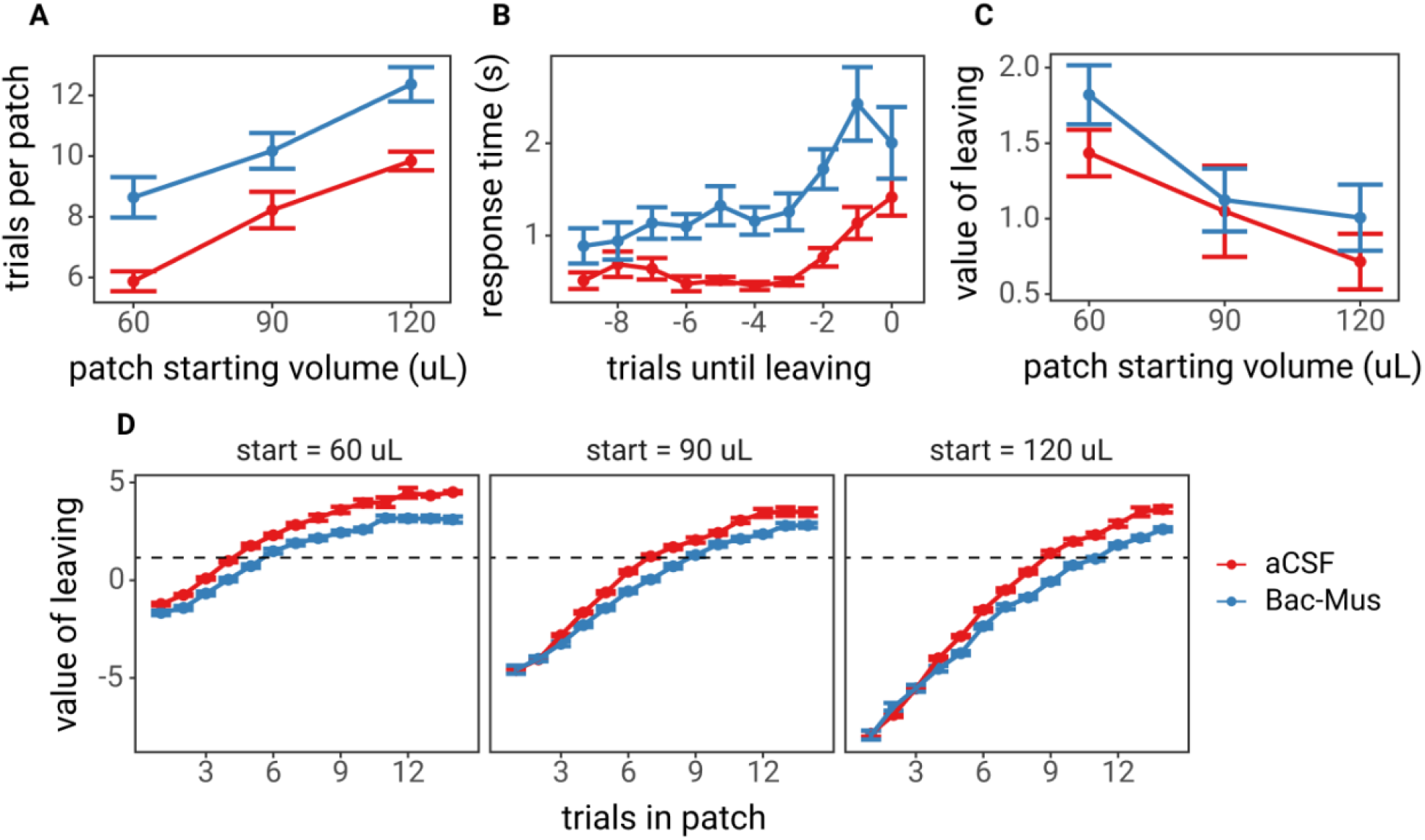
Effects of Inactivation of ACC on foraging behavior. A) The number of trials spent in each of the three patch types. B) Response times as rats became closer to leaving patches, where 0 trials until leaving is the nose poke to leave the patch, -1 is the last lever press in the patch, and so on. C) The value of leaving (global reward rate – local reward rate) the rat experienced on the last trial in which they harvested reward from the patch (-1 trials until leaving). D) The value of leaving over the course of trials in the patch for each of the three patch starting rewards. The horizontal dashed line represents the average value of leaving at which rats chose to leave the patch. The intersection of the plotted value of leaving (red and blue lines) and average value of leaving is where rats would be expected to leave the patch. In all plots, points and lines represent the mean of means for each rat, and error bars represent the standard error of the mean across rats.

That rats stay longer in patches and exhibit longer response times due to ACC inactivation suggests that ACC is involved in the foraging decision process. However, MVT predicts that foraging decisions are based on estimates of reward rate – not reward value – and animals that exhibit longer response times experience lower reward rates. Thus, staying longer in patches could be a compensatory mechanism for lower reward rates experienced as a consequence of longer response times. The relative value of leaving, the difference between the global and local reward rate, over the course of trials in patches was lower during sessions in which rats were injected with Bac-Mus compared to aCSF (Figure 6D). Furthermore, there was little to no difference in the value of leaving on the last decision to stay in patches between Bac-Mus and aCSF sessions (main effect of Bac-Mus vs. aCSF: *β* = 0.245, SE = 0.128, F(1, 887) = 3.690, p = .055; Figure 6C). Finally, to determine whether ACC inactivation may have caused rats to move slower, we examined the time it took rats to move from the lever to the reward magazine, a period of time that reflects only movement as no decision needs to be made. Rats were slower to move from the lever to the reward port during Bac-Mus sessions compared to aCSF sessions (t(10) = 5.649, p < .001, paired t-test; Figure 6-1A). These findings support the notion that ACC inactivation may have slowed response times unrelated to decision deliberation, causing rats to stay longer in patches to compensate for lower reward rates.

To better understand how ACC contributes to the foraging decision process, the LCA model was fit to rat behavior on aCSF and Bac-Mus sessions, and we compared the difference in the best fit parameters from these different sessions. Model predictions for aCSF and Bac-Mus sessions, and all parameter estimates are shown in Figure 6-1. The only parameter that was significantly different across sessions (paired t-tests, p < .05, holm correction for multiple comparisons across 13 parameters; Figure 6-2) was the non-decision time, *η*. This finding further supports the hypothesis that changes in foraging decisions and response times due to ACC inactivation were not related to altered encoding of the value of staying vs. leaving, nor to changes in the accumulation to bound decision process. ACC is more likely serving a more general function such as performance monitoring and regulation of cognitive control. Alternatively, longer response times may reflect a reduction in motivation or response vigor. Although ACC is not directly involved in a value comparison process, the ACC may play an important role in regulating vigor as a function of the global and/or local reward rates.

## Discussion

Previous studies have shown that ACC activity is modulated during foraging decisions (Hayden et al., 2011; Kolling et al., 2012, 2014; Blanchard and Hayden, 2014; Shenhav et al., 2014, 2016b), but what this modulation of ACC activity represents, or how the ACC contributes to foraging decisions, is not fully understood. In the present study, we found that rat ACC neurons separately correlate with both foraging decisions and response times. Using a LCA model that estimates MVT-derived decision variables within and across trials, we found that individual ACC neurons encode lower-level task variables such as the reward rate and decision accumulators, and that as a population, the average ACC activity reflects decision difficulty as indexed by the similarity in the value of staying in the patch vs. leaving. Finally, inactivation of ACC neurons altered foraging behavior, but this change was best explained not by changes to the decision process, but by altering non-decision time, a latent variable meant to represent the time for non-decisional sensory processing and motor execution, which may include motivation or response vigor.

The findings that ACC neurons continuously signal important foraging decision variables, but that the ACC is not necessary to follow the MVT decision rule, may provide important information about its function. Despite encoding value signals that could be used for decisions, results from the ACC inactivation experiment indicate that the ACC is not necessary for comparing the values of options for the decision at hand in the task we employed. The lack of involvement of ACC in the primary decision strategy is consistent with a recent study that performed optogenetic silencing of ACC in mice performing a foraging-style task (Vertechi et al., 2020). In this study, mice chose to stay vs. leave a patch for which the probability of receiving a reward was reduced with every decision to stay, and decisions to stay vs. leave were equally guided by failures to receive reward despite ACC inactivation. Ultimately, these results support a more general role for the ACC in monitoring peformance and regulating cognitive control during foraging tasks, a function long associated with the ACC (Botvinick et al., 2001), and a role that ACC has been previously hypothesized to perform in foraging tasks (Blanchard and Hayden, 2014; Shenhav et al., 2014, 2016b). Another possible interpretation of the effect of ACC inactivation on non-decision times is that the ACC may play a role in setting response vigor, or the speed at which animals choose to perform a task. Response vigor is thought to be an integral aspect of foraging decisions – when the global reward rate is higher, it is beneficial for animals to exhibit greater vigor in order to reap the benefits of a more rewarding environment (Niv et al., 2007). While the ACC may not contribute to setting the patch leaving threshold, it may play a critical role in setting response vigor based on the estimated patch-leaving threshold.

One critique of the hypothesis that ACC serves as a monitor of performance is that although performance monitoring signals are often seen in average ACC activity measured using fMRI in humans, performance monitoring variables such as decision difficulty or response competition have not been found in ACC in many single unit studies (Heilbronner and Hayden, 2016; Kolling et al., 2016). A model to illustrate this concept has been proposed previously by Kolling et al., 2016. In this report, PCA analyses revealed that although a performance monitoring variable, decision difficulty, is evident in average ACC activity, the components of that average ACC signal (single- and multi-units) vary more closely with lower-level decision variables. This analysis, combined with results from inactivation studies, illustrates that encoding of lower-level decision variables, such as reward rates and decision accumulators, are components of a more general system that regulates cognitive control or motivation.

Furthermore, it is possible that ACC does not play a critical role in foraging decisions that do not involve model updates, but that the ACC will become critical if there is a change in the foraging environment (e.g. a change in the possible patch types that animals may encounter. Previous studies have found that perturbation of ACC activity affects animals’ ability to perform task or strategy switching or to update internal models of the environment (Kennerley et al., 2006; Tervo et al., 2014; Sarafyazd and Jazayeri, 2019; Akam et al., 2020). This kind of model updating does not apply to traditional foraging tasks, such as the one used here. In foraging tasks, even if there is some uncertainty about the exact reward to be received in future patches (e.g. if there are multiple patch types), animals have learned the average expected future reward. Thus, an animal can learn the appropriate time to leave different patch types without updating internal models of the environment.

These findings suggest that rat ACC activity correlates closely with decision difficulty in a foraging task, similar to foraging-related ACC activity that has been reported in humans and non-human primates (Hayden et al., 2011; Shenhav et al., 2014, 2016b). Although the degree of homology between rodent and primate cingulate cortex has not been entirely clear (Seamans et al., 2008; Heilbronner and Hayden, 2016; Heilbronner et al., 2016; van Heukelum et al., 2020), these findings contribute to a growing body of literature suggesting that rodent ACC exhibits similar activity to human and non-human primate ACC. Recent studies that have found that rodent ACC neurons exhibit signals that have long been associated with human ACC, including feedback-related negativity (Warren et al., 2015), error monitoring (Narayanan et al., 2013), and increased activity during response competition (Bryden et al., 2018). Furthermore, a recent lesion study found that the ACC is necessary to resolve response competition (Brockett et al., 2020). Together, these studies present a case that rodents can serve as a model to understand the function of the ACC and the behavioral consequences of ACC dysfunction.

## Acknowledgements

Author contributions: All authors participated in study design; GAK, MHJ collected data; GAK, AS analyzed data; All authors wrote the paper.

This work was supported by NIH grants F31MH109286 (GAK), R01MH124849 (AS), R01MH092868 (GAJ). The authors would like to thank Jeremy Autore for assistance with computational modeling, and the Shenhav Lab at Brown University for valuable discussions on this work.

## Extended Data

**Figure 2-1.**
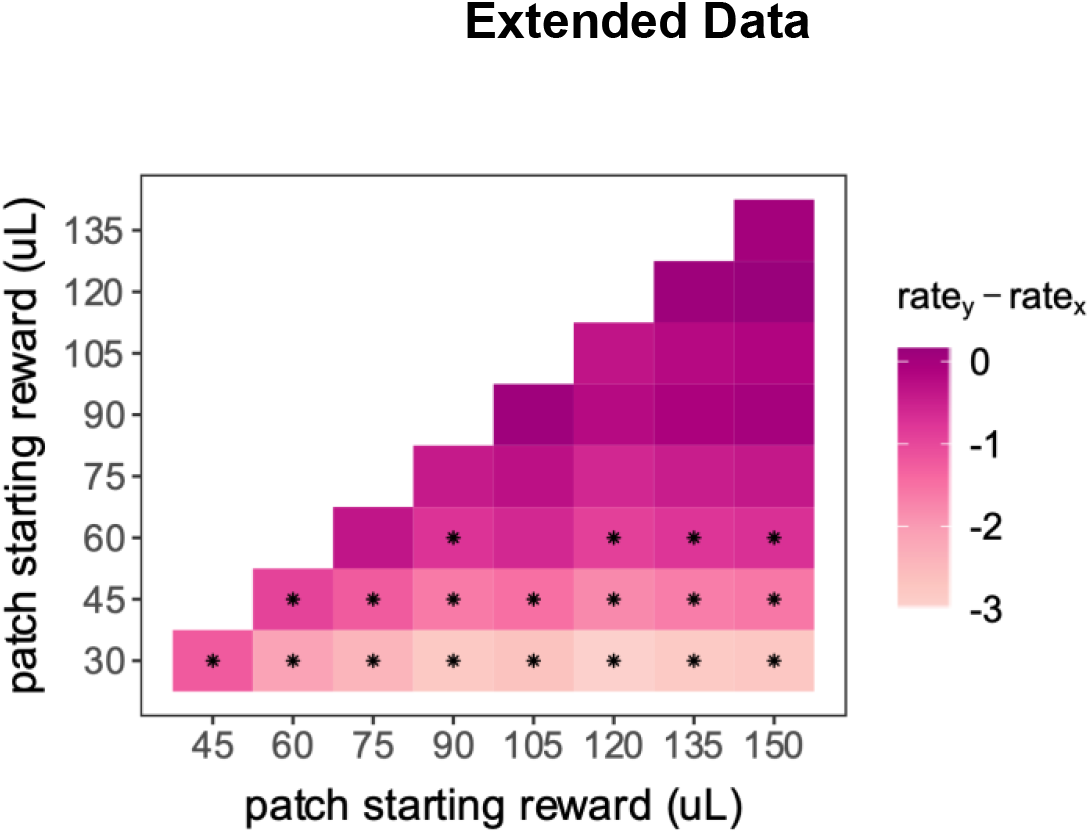
Pairwise comparisons of the patch leaving threshold (the local reward rate when rats decided to leave the patch), for all patch types. Colors indicate the difference in the reward rate threshold at which rats decided to leave the patches (i.e., y-axis leaving threshold – x-axis leaving threshold). Asterisks indicate statistically significant comparisons (pairwise chi-square tests with holm correction).

**Figure 3-1.**
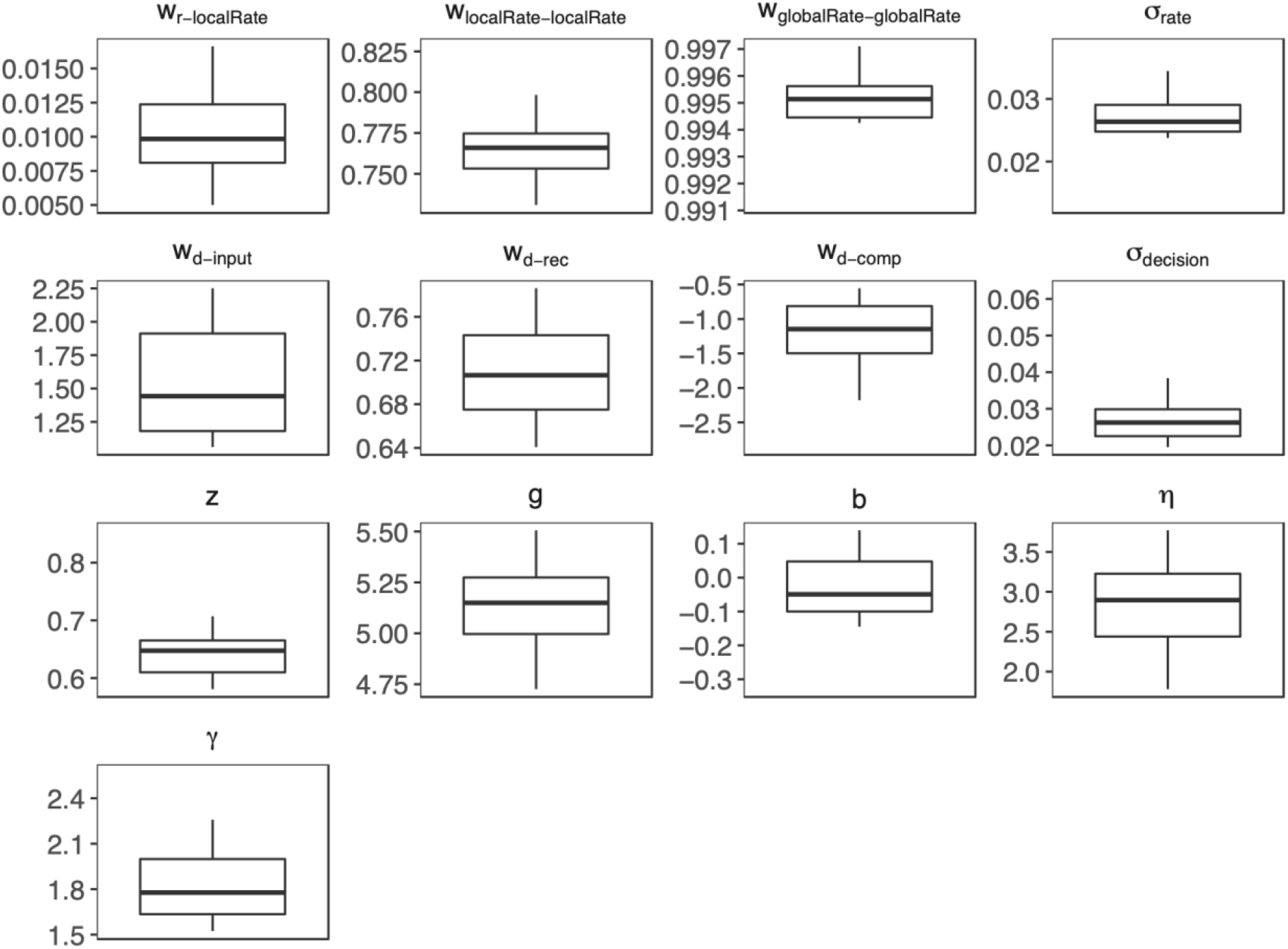
Parameter estimates for LCA fits to animal behavior during recording sessions. Each panel shows the distribution of parameter estimates across animals for a given parameter. The box represents the first and third quartile, whiskers represent 1.5 times the interquartile range, with all individuals plotted transparently.

**Figure 3-2.**
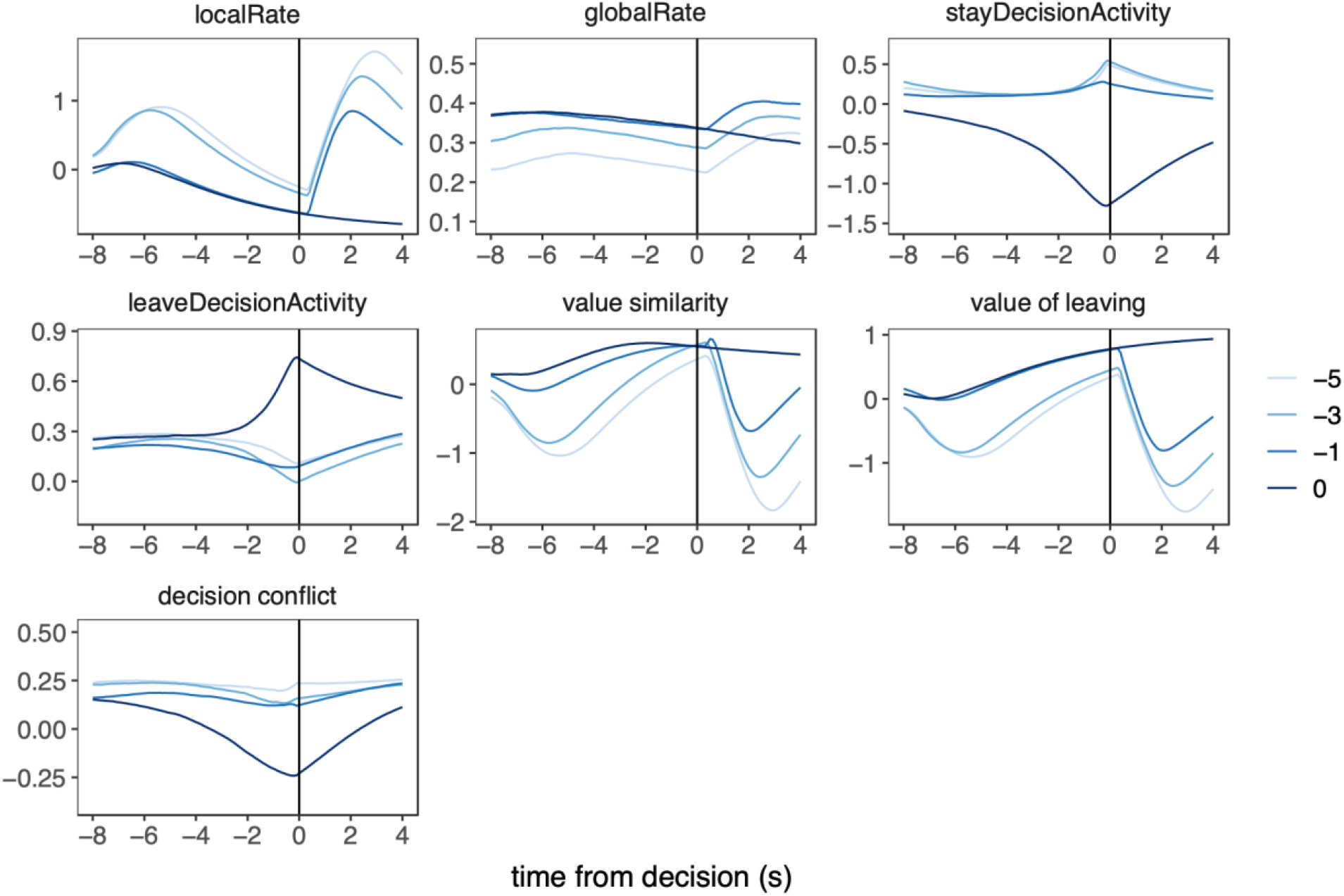
PETHs of LCA model units and hypotheses for ACC activity in the foraging task. Each panel represents the PETH of activity of LCA model units (*localRate, globalRate, stayDecisionActivity, leaveDecisionActivity*) or hypotheses for ACC activity measured from LCA model units (value similarity, value of leaving, and decision conflict) during a simulation of the foraging task. For each PETH, time = 0 represents the time of the lever press to harvest reward or nose poke to leave the patch, and colors represent the number of trials until leaving the patch.

**Figure 4-1.**
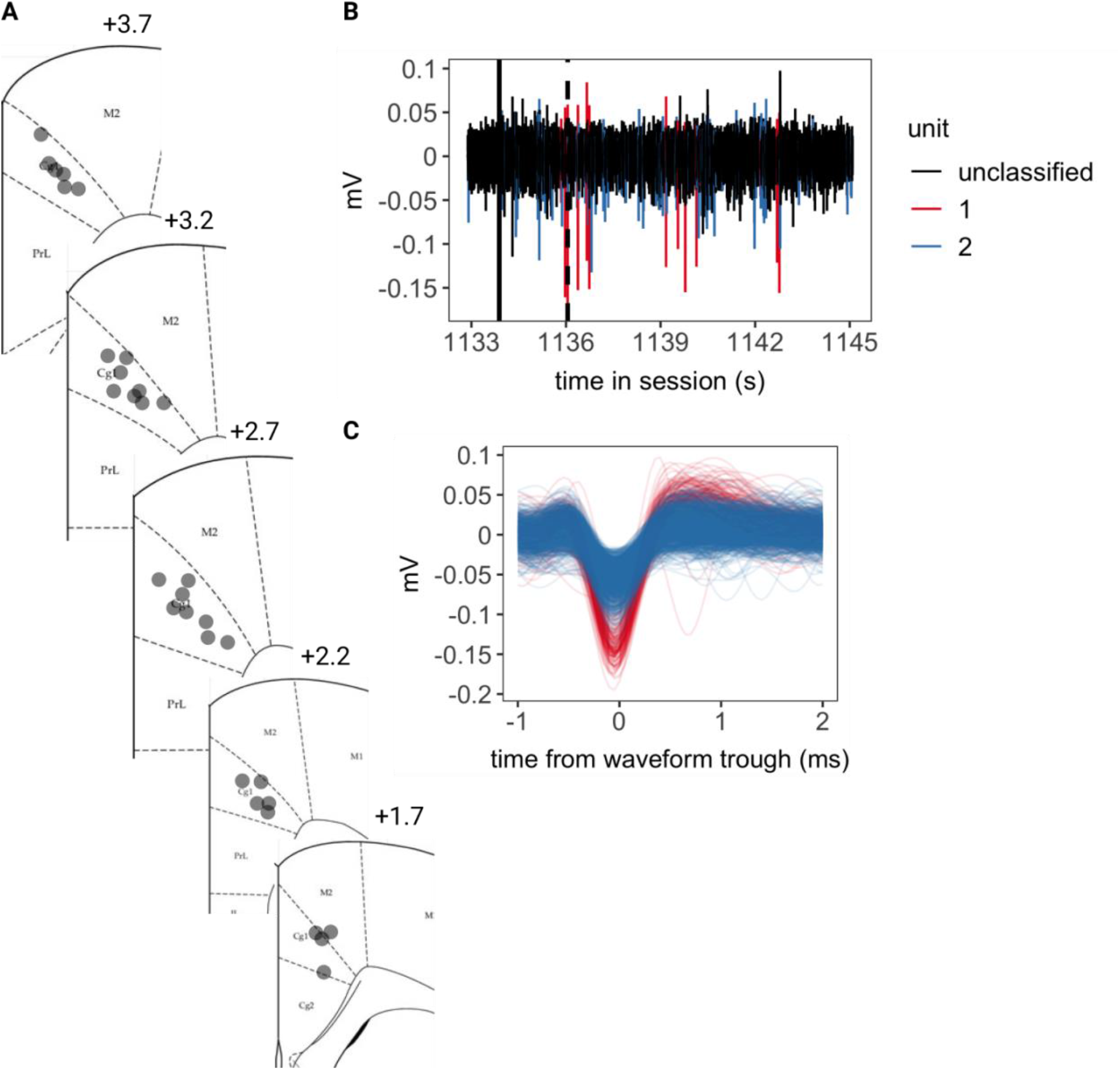
A) Diagram of identified recording locations localized in the Cg1 region. B) An example trace of a bandpass filtered signal (300-3000 Hz) of a single channel for the duration of one trial. The solid black line indicates the start of the trial, and the dashed black line indicates the time of the lever press. The colored portion of the trace indicates spikes assigned to one of two units on this channel. C) The waveforms of the units in B.

**Figure 4-2.**
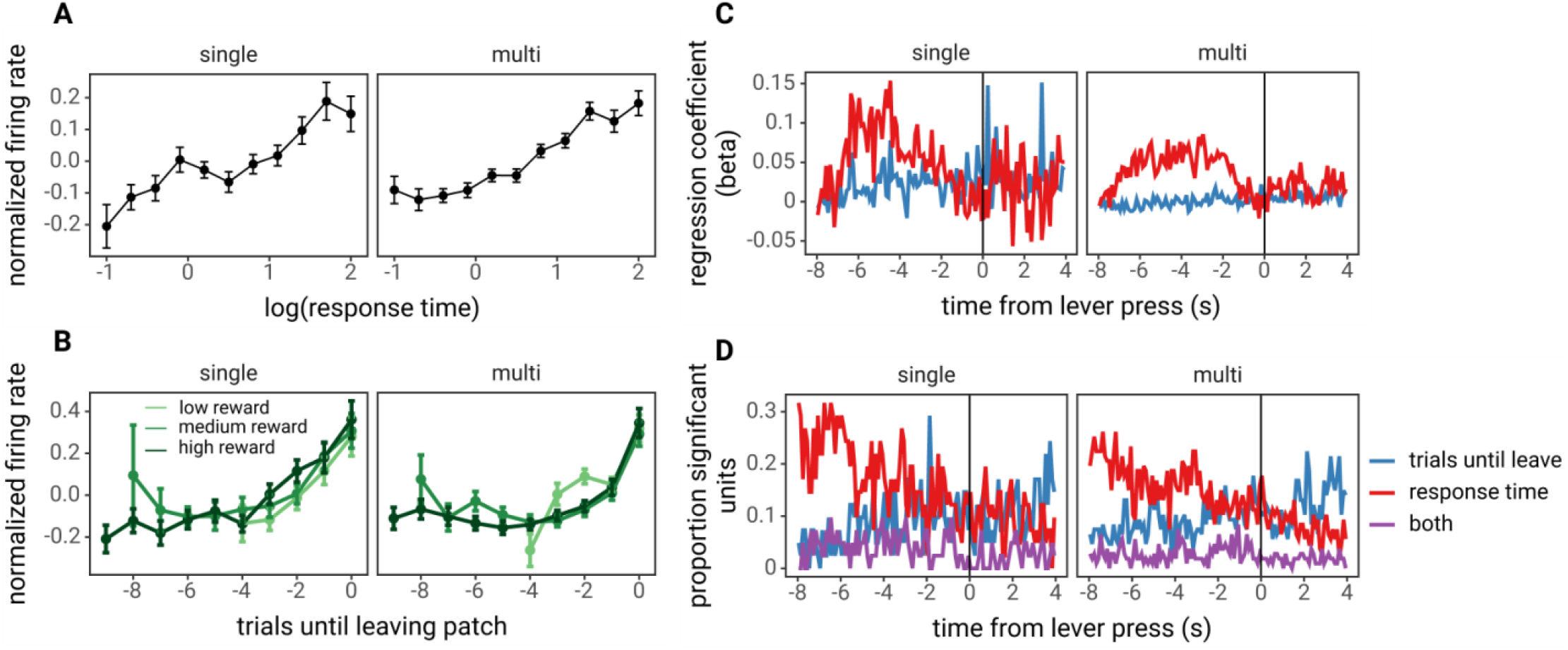
Comparison of activity and encoding of single- vs. multi-units. A) The average normalized firing rate (z-scored) as a function of the log of the response time (as in Figure 4B). B) Average normalized firing rate as a function of the trials until leaving the patch. Colors indicate patches ACC activity in patches start with low, medium, and high reward. C) The average effect of trials until leaving and response times at each time point within a trial, locked to the time of the lever press to saty in the patch. D) The proportion of untis with significant effects of trials until leaving, response times, or both (P < .05, z-test on regression coefficient) at each time point within the trial.

**Figure 5-1.**
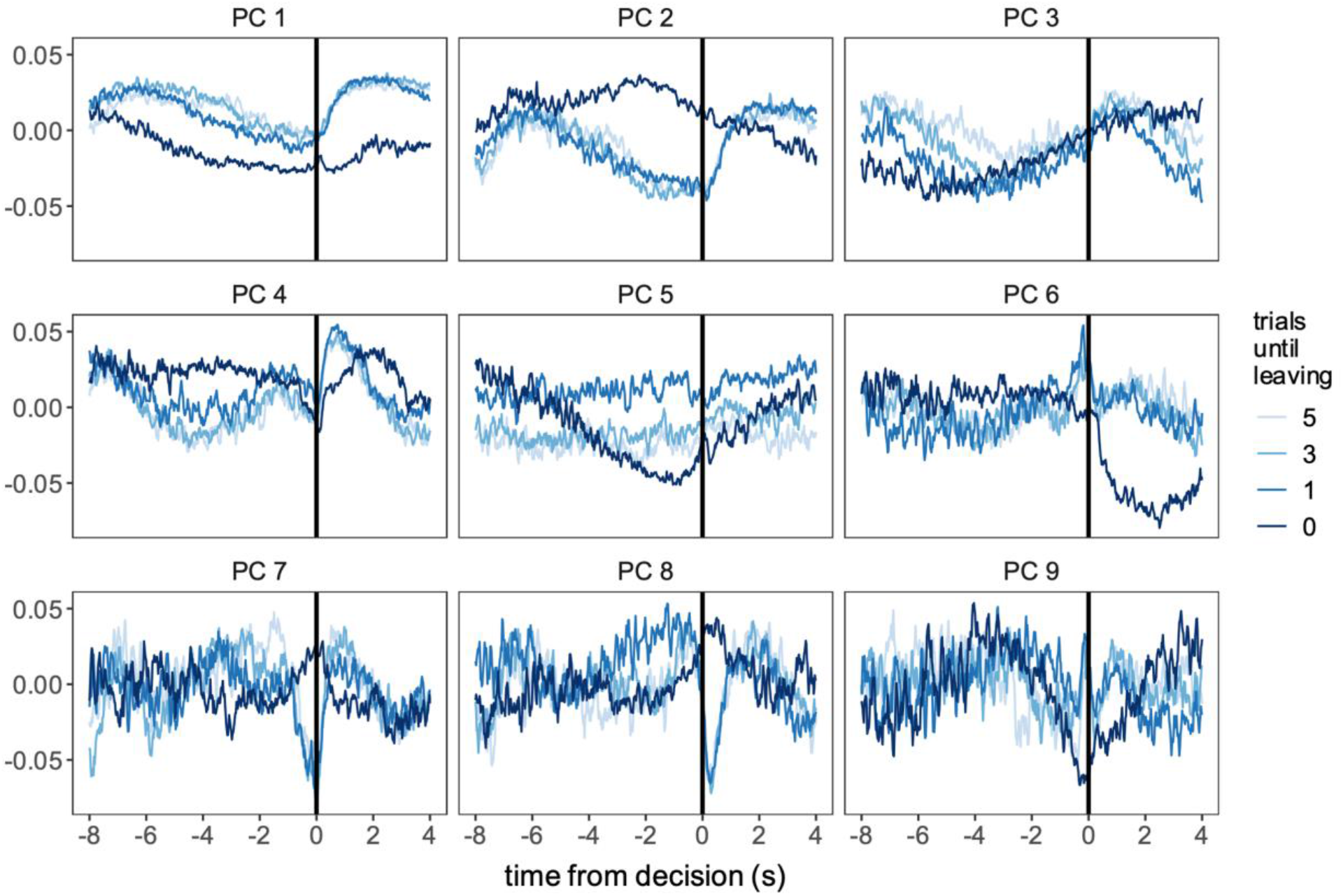
The first 10 principal components across all individual units for 5, 3, 1, and 0 trials until leaving the patch.

**Figure 6-1.**
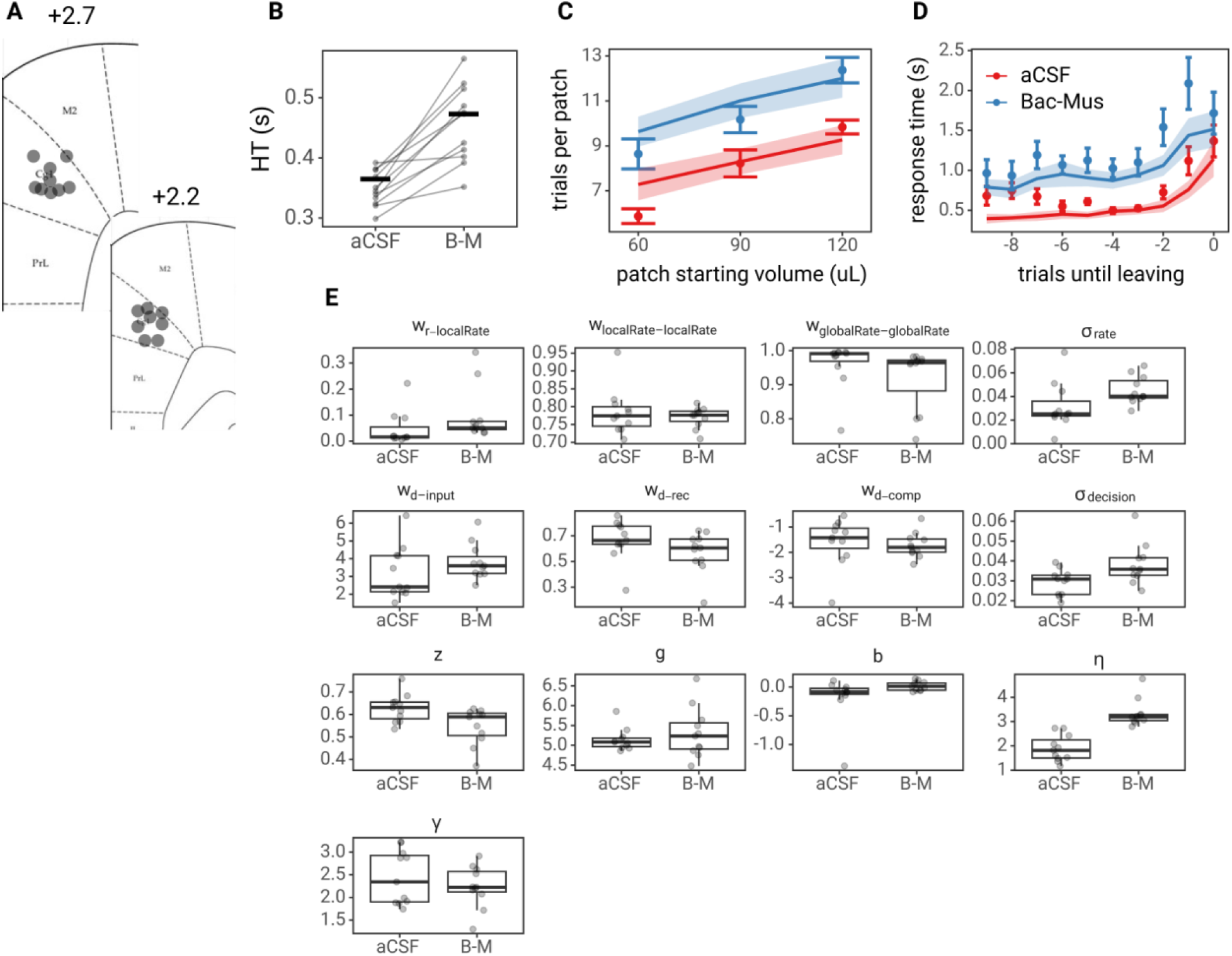
LCA model predictions for the ACC inactivation experiment. A) Diagram of identified cannula placements for the ACC inactivation experiment. Numbers above diagrams indicate the distance from bregma. B) The handling time (HT) or time from lever press to entering the reward port on aCSF and Bac-Mus (B-M) sessions. Grey lines indicate data for each individual animal and black, horizontal dash indicates the mean for each treatment. C) Predicted number of trials spent in patches plotted against observed rat behavior. D) Predicted response times against observed rat behavior. Points and error bars represent the mean and standard error of rat behavior, and lines and ribbon represent the mean and standard error of model predicted behavior. E) LCA model parameters fit to aCSF and Bac-Mus (B-M) sessions. The box represents the first and third quartile, whiskers represent 1.5 times the interquartile range, with all individuals plotted transparently.

**Figure 6-2.**
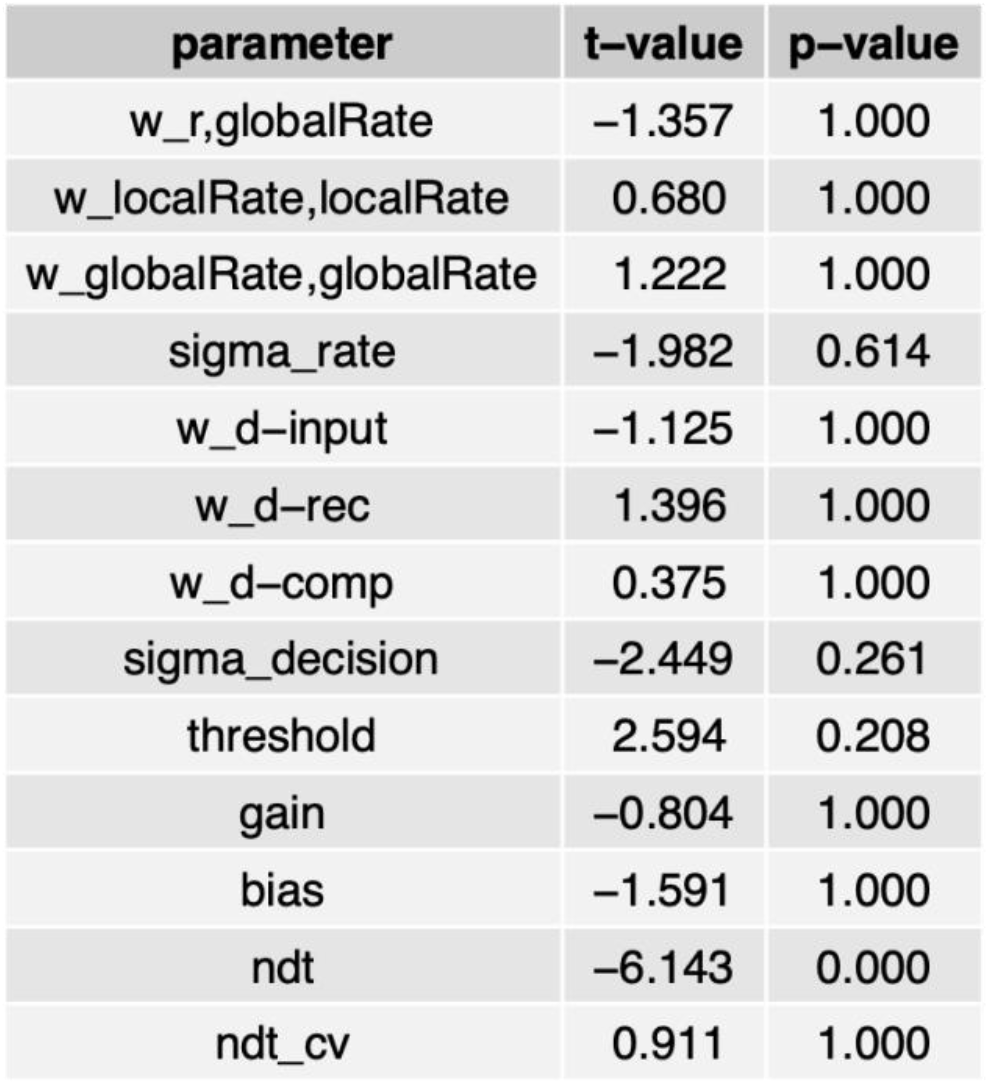
Pairwise t-tests on LCA model parameters fit to aCSF sessions vs. parameters fit to Bac-Mus sessions.

